# Open source 3D phenotyping of chickpea plant architecture across plant development

**DOI:** 10.1101/2020.09.08.288530

**Authors:** William T. Salter, Arjina Shrestha, Margaret M Barbour

## Abstract

In this work, we developed a low-cost 3D scanner and used an open source data processing pipeline to phenotype the 3D structure of individual chickpea plants. Being able to accurately assess the 3D architecture of plant canopies can allow us to better estimate plant productivity and improve our understanding of underlying plant processes. This is especially true if we can monitor these traits across plant development. Photogrammetry techniques, such as structure from motion, have been shown to provide accurate 3D reconstructions of monocot crop species such as wheat and rice, yet there has been little success reconstructing crop species with smaller leaves and more complex branching architectures, such as chickpea. The imaging system we developed consists of a user programmable turntable and three cameras that automatically captures 120 images of each plant and offloads these to a computer for processing. The capture process takes 5-10 minutes for each plant and the majority of the reconstruction process on a Windows PC is automated. Plant height and total plant surface area were validated against “ground truth” measurements, producing R^2^ > 0.99 and a mean absolute percentage error < 10%. We demonstrate the ability to assess several important architectural traits, including canopy volume and projected area, and estimate relative growth rate in commercial chickpea cultivars and lines from local and international breeding collections. Detailed analysis of individual reconstructions also allowed us to investigate partitioning of plant surface area, and by proxy plant biomass.

## Introduction

With a growing global population and increasingly challenging environmental conditions, it is critical for plant scientists to think ‘outside the box’ to identify novel plant traits that could be used to improve crop yield. This could be through the adoption of new technologies, through investigation of phenotypic traits throughout plant development or through improvement of crops that have been bred less intensively in the past. Recent advances in image capture and computing technologies have allowed us to accurately phenotype the 3D architecture of crop plants such as wheat, barley and rice. However, structurally complex crop species, such as chickpea, remain elusive due to their small leaves, high levels of branching and indeterminant nature. Here we assess the potential to phenotype the 3D structure of these “complex” plants across their development using photogrammetry.

One of the main problems with conventional, direct measurements of plant structural properties is that they are laborious and often destructive. This is particularly evident when working with larger plants and plant species with many small leaves. Imaging intact plants can bypass the need to destructively harvest plants, allowing for the measurement of structural traits across plant development. Two-dimensional imaging techniques have long been used for the quantitative measurement of plant structural traits, including plant surface area, number of leaves, leaf shape and leaf colour (a database of such approaches is presented in Lobet et al, 2013; and is continually updated). Using new software tools, such as PlantCV (Gehan et al., 2017), quantitative traits can even be extracted automatically from images, reducing user error and analysis time. However, most 2D imaging techniques were developed to only work for small plants with a simple structure, such as the two-dimensional rosettes of the model plant *Arabidopsis thaliana* (Vasseur et al., 2018), or to only extract relatively basic information, such as plant height (Atieno et al., 2017). For more complex or larger plants, 2D imaging techniques can result in inaccuracies due to overlapping features in captured images (i.e. occlusion of stems by leaves, leaves by leaves, etc.). 3D imaging addresses this issue, allowing us to capture the full detail of a plant’s structure without self-occlusion of any plant tissues.

There are several methods available to phenotype the 3D structure of plants (for detailed reviews see Gibbs et al., 2017 and Paulus, 2019). Laser scanning (LiDAR) can provide very detailed reconstructions of plants but there is often a trade-off between the cost of instrumentation and the complexity of 3D models. Commercial instruments can cost upwards of US $10,000 but can generate detailed models of plants with > 2 million points. Newly developed DIY instruments can cost as little as US $400 but only generate models with approx. 40,000 points (Panjvani et al., 2019). LiDAR can also be inflexible, both in terms of sample size (i.e. one system may provide good resolution for small plants but not for large plants, and vice versa) and downstream data analyses (i.e. may be limited to certain commercial data analysis programs). Photogrammetry on the other hand can be highly cost effective and versatile. Photographs of the plant are taken from multiple angles using a standard camera and subsequent computer analyses is used to reconstruct a scaled 3D model. This 3D reconstruction can then be used for trait measurements, such as plant dimensions, plant surface area and leaf area index, and modelling simulations, such as ray tracing of the canopy light environment (Pound et al., 2014). Data quality can be comparable to more expensive LiDAR systems and it can be used for subjects of wide-ranging sizes. Generally speaking, the more photos of the subject, the better the reconstruction will be with regards to precision and accuracy (Nguyen et al., 2016), albeit with a longer capture time. Many photogrammetry software packages are open source (including Colmap, Schonberger & Frahm, 2016; Meshroom, Alicevision, 2018; and VisualSFM, Wu, 2011), meaning that they are freely available and can be modified at the code level to give users a highly customised and powerful experience. Photogrammetry has been used effectively for the 3D reconstruction of a number of monocot crop species, including wheat (Burgess et al., 2015) and rice (Burgess et al., 2017), and for species with larger leaves, such as sunflower (Gélard et al., 2017) and soybean (Zhu et al., 2020). However, few studies have assessed whether it could be used to accurately reconstruct 3D models of plant species with numerous small leaves, such as chickpea.

Chickpea (*Cicer arietinum* L.) has long been an important annual crop for resource poor farmers across the globe but there is growing demand elsewhere due to changing diets and a push for protein rich alternatives to meat (Thudi et al., 2016). Chickpea is often considered more sustainable than non-legume grain crops, such as wheat or rice, due to its ability to form symbiotic relationships with nitrogen fixing bacteria, reducing reliance on nitrogen fertiliser (Downie, 2014). It can also be used effectively in rotation with cereal crops to break the life cycle of diseases and improve soil health (Marcellos et al., 1998). Chickpea can therefore be a lucrative option for many growers, particularly considering there are also economic benefits, with returns to Australian growers of roughly AU $300 t^−1^ compared to around AU $100 t^−1^ for wheat between 2012 and 2014 (GRDC, 2017). Yet, whilst chickpea has an estimated yield potential of 6 t ha^−1^ under optimal growing conditions, annual productivity of chickpea worldwide currently sits at less than 1 t ha^−1^ (Thudi et al., 2016). This yield gap is the result of a lack of genetic diversity in breeding programs that has left cultivars susceptible to biotic and abiotic stresses. Phenotyping for natural variation in traits of interest across diverse germplasm could be used to minimise this yield gap and to improve grain yield potential. Chickpea is an indeterminate crop in which vegetative growth continues after flowering begins. This growth form poses management challenges for growers (William & Saxena, 1991) and can result in yield losses. issues. Growers may need to strategically terminate vegetative growth using herbicide before harvest, posing health issues for the farmer and the environment. There has been a report of determinacy in chickpea (Hedge, 2011), as well as the identification of regulatory genes in other bushy legume species including common bean (Kwak et al., 2012), pigeonpea (Saxena et al., 2017) and soybean (Tian et al., 2010). These reports highlight that determinacy is a genetically controlled trait that could be explored by phenotyping the growth habit of diverse chickpea populations. Chickpea also has a highly branching structure, requiring more resources to be allocated to structural tissue, which may reduce remobilisation of nutrients to pods during reproductive growth (Alerding et al., 2018). Modification of plant architecture through targeted plant breeding has led to huge successes in other crop species, most notable was the introduction of dwarfing genes into elite varieties of wheat, which led to increased seed yields, reduced yield losses due to lodging and was integral to the green revolution of the 1960s and 1970s (Jobson et al., 2019). By phenotyping canopy architecture traits across chickpea genotypes, we will improve our understanding of the underlying genetics controlling these traits, how these traits influence plant productivity and can then use this information to make informed breeding decisions.

The main aim of this work was to develop and validate a low-cost and open source photogrammetric method for detailed 3D phenotyping of chickpea plants. The imaging setup consisted of three DSLR cameras, LED lighting and a motorised turntable, controlled by a user-programmable Arduino microcontroller. 3D reconstruction and analyses of 3D models were performed using open-source software on a Windows PC. The system was tested with a variety of chickpea genotypes (three commercial and three pre-breeding lines) and phenotyping measurements were validated against conventional, destructive measurement techniques. We also assessed whether differences in plant architecture or growth rates could be observed across chickpea genotypes.

## Methods

### Plant material

Three commercial Australian chickpea (*Cicer arietinum* L.) cultivars (PBA Slasher, PBA Hattrick and Genesis Kalkee) were grown from seed in a controlled glasshouse in August 2019. These genotypes were selected as their architecture is known to differ in the field (Table S1) and are referred to collectively herein as “commercial cultivars”. Seeds were planted in potting mix containing slow release fertiliser (Osmocote Premium; Evergreen Garden Care Australia, Bella Vista, NSW, Australia) in 7 l square pots and watered to field capacity once daily. The daytime temperature in the glasshouse was controlled to 25°C and the nighttime temperature controlled to 18°C. The relative humidity was set to 60%. Supplemental lighting was provided by LED growth lights if ambient light fell below a photosynthetic photon flux density (PPFD) of 400 µmol m^−2^ s^−1^, this effectively maintained a PPFD of > 400 µmol m^−2^ s^−1^ at the plant level at all times during the day. Fifteen plants (five for each genotype) were transferred from the glasshouse to the laboratory for imaging once per week and were returned to the glasshouse after measurement. Additionally, each week fifteen plants (five of each genotype) were imaged and then destructively harvested for validation of 3D scanner measurements.

Three chickpea genotypes (ICC5878, SonSla and PUSA76) were selected from local and international sources based on contrasting canopy architecture and growth-related traits (Table S1) and are referred to collectively herein as “breeding lines”. ICC 5878 is from the ICRISAT Chickpea Reference Set (http://www.icrisat.org/what-we-do/crops/ChickPea/Chickpea_Reference1.htm). SonSla is a fixed line (F7-derived) resulting from a cross between Australian cultivars Sonali and PBA Slasher. PUSA 76 is an older accession released by IARI, India and imported via the Australian Grains Genebank. These were grown outside in February – April 2020. Seeds were planted in potting mix containing slow release fertiliser (Complete Vegetable and Seedling Mix; Australian Native Landscapes Pty Ltd, North Ryde, NSW, Australia) in 7 l square pots and watered every three days to field capacity. Twelve plants of each genotype were imaged at five weeks post-germination and destructively harvested for validation of 3D scanner measurements.

### Semi-automated 3D imaging platform

Plants were imaged using a turntable and camera photogrammetry setup (schematic in Figure 1). The turntable is constructed from acrylic (Suntuf 1010493; Palram Australia, Derrimut, Victoria). It consists of a circular top plate on which the potted plant is placed and a base which houses a stepper motor (42BYG; Makeblock Co., Ltd, Shenzhen, China). A lazy susan bearing plate (Adoored 0080820; Bunnings Warehouse, Hawthorn East, Victoria, Australia) is used to connect the plate to the base to provide smoother movement and reduce strain on the motor during imaging. The turntable is connected to and controlled by a user-programmable Arduino microcontroller (Uno R3; Arduino LLC, Somerville, MA, USA) and a number of Arduino breakout boards. The stepper motor is driven via a stepper driver board (DRV8825; Pololu, Las Vegas, NV, USA), that provides precise control of turntable rotation, allowing for individual rotational microsteps as small as 0.06°. A copper heatsink (FIT0367; DFRobot, Shanghai, China) and 5 V fan (ADA3368, Adafruit Industries LLC, New York, NY, USA) are installed on the stepper driver to prevent overheating. The microcontroller triggers the cameras via a relay breakout board (Grove; Seeed Studio, Shenzhen, China) and a custom-made remote shutter cable. An LCD screen with integrated keypad (DFR0009; DFRobot) is used to operate the turntable and provide basic information during the capture process. A 5 V buzzer (AB3462; Jaycar, Sydney, NSW, Australia) audibly alerts the user when a full rotation is complete. Power is provided via a mains – 12 V DC 5 A power supply (MP3243; Jaycar). The motor is powered directly with 12 V DC whilst a step-down voltage regulator (XC4514; Jaycar) is used to provide 5V DC to the microcontroller and associated boards. A wiring diagram is provided in Figure 1b. The turntable was set on a white table against a white backdrop (Figure 2).

**Figure 1.**
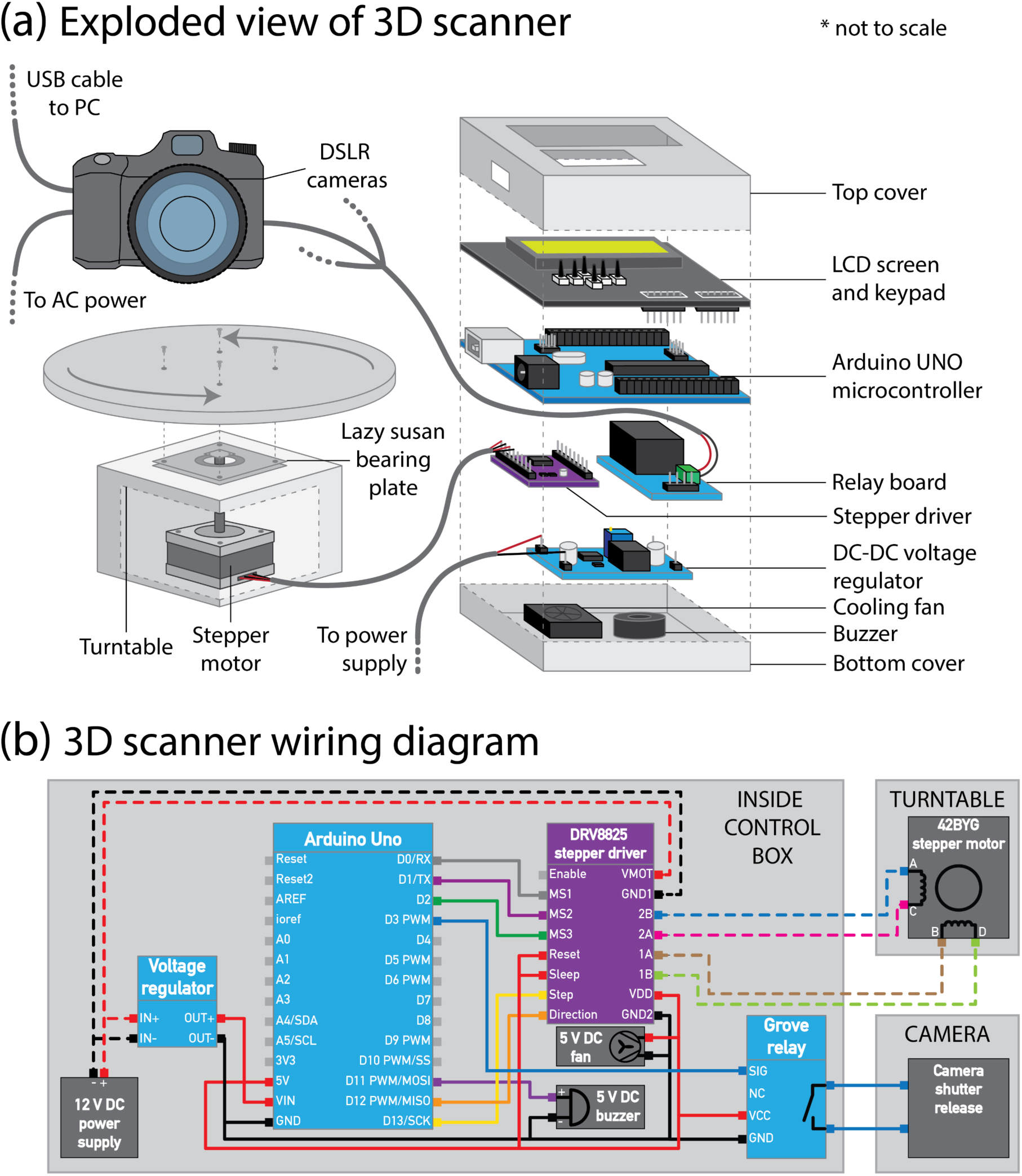
**(a)** Exploded view of the 3D scanner showing components required for assembly. (b) Wiring diagram for the 3D scanner. Note that in **(b)**, 5 V wires are represented by solid lines and 12 V wires by dashed lines.

**Figure 2.**
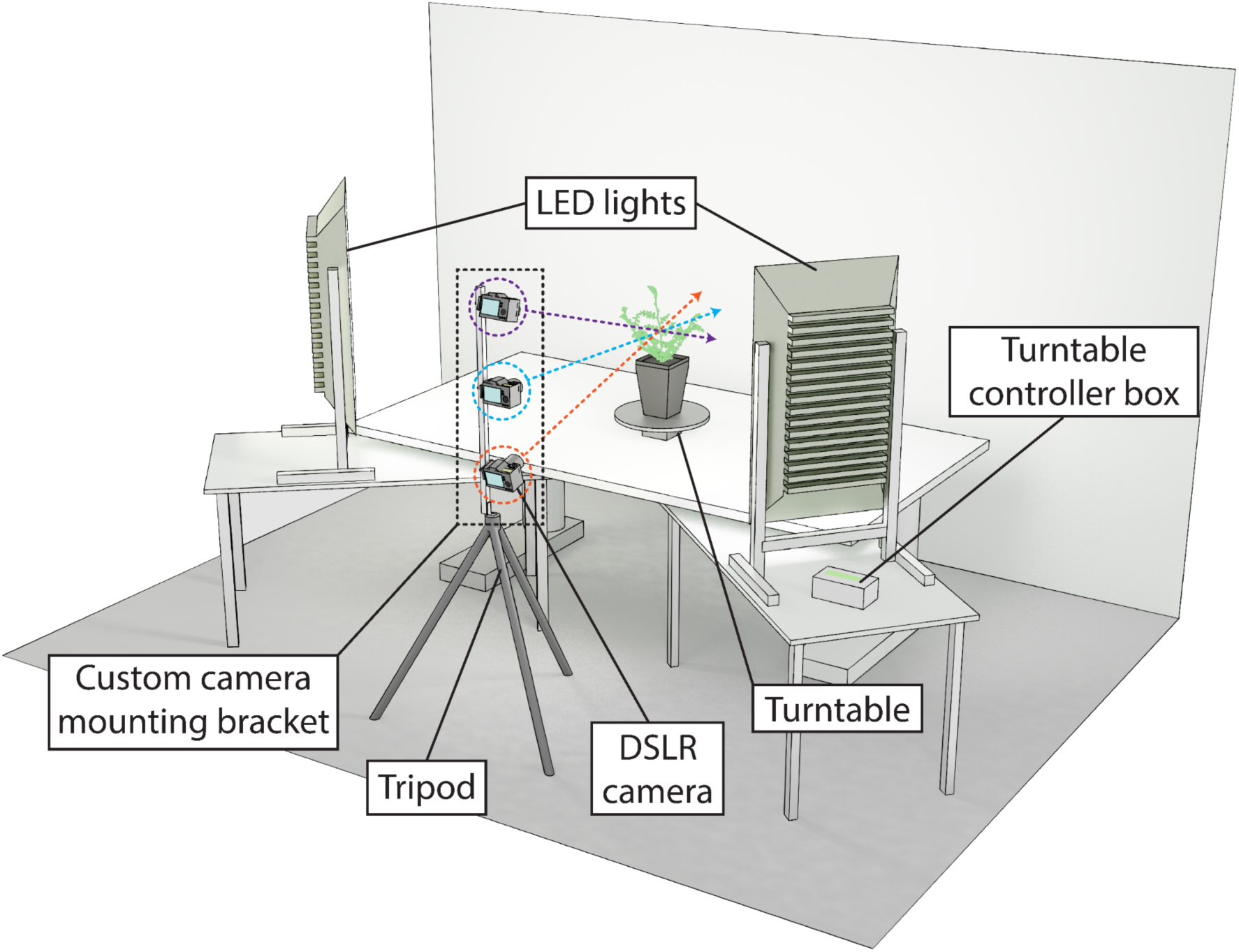
Diagram showing the 3D scanner set-up in the laboratory. The coloured circles highlight the three cameras angled to face the plant. Note that no cables are shown in the diagram for the purpose of clarity. Exact spacing of the set-up is shown in Figure S1.

The microcontroller is programmed using the open source Arduino IDE software (Version 1.8.10; Arduino LLC). The automated capture program was designed such that it will turn the plant a set number of degrees (determined by the user), pause briefly for the plant to stop moving (with a delay programmed by the user) and then trigger the camera(s) to capture an image. This process is repeated until a full rotation of the plant has been captured. The microcontroller also offers the user some control of the turntable via the buttons on the LCD shield (to increase/decrease the number of images captured per rotation, to manually turn the plant clockwise/anticlockwise and to start/pause/stop the automated capture sequence). Further control of the capture sequence can be achieved through modification of the code. The Arduino program is provided as a supplementary file.

Lighting is provided by two large LED floodlights (generic LED floodlights bought on eBay) held in a vertical orientation with custom stands made from aluminium extrusion (Figure 2a). A sheet of white acrylic (Suntuf 1010493; Palram Australia) is placed over the front of each light as a diffuser. Large cooling fans (MEC0381V3; Sunon, Kaohsiung City, Taiwan) were installed on the rear of the lights. In our imaging setup, the lights were set 80 cm away from the plant on either side of the tripod and angled to face the plant directly (Figure S1).

A tripod (190XPRO; Manfrotto, Cassola, Italy) was used as a base for a custom-made camera mounting bracket (schematic in Figure S2). The top of the tripod was set level with the table on which the turntable was sat. The mounting bracket was constructed from a 110 cm length of aluminium square hollow extrusion with three quick release mounting points (323 Quick Change Plate Adapter; Manfrotto) for a camera positioned at 10 cm, 55 cm and 100 cm vertically from the base and angled towards the plant. A steel angle bracket (SAZ15; Carinya, Melbourne, Australia) was bolted to the bottom of the aluminium extrusion for secure attachment to the tripod.

### Camera setups

Three digital SLRs (D3300; Nikon Corporation, Tokyo, Japan) were used for imaging, each with a 50 mm prime lens (YN50; Yongnuo, Shenzhen, China). The cameras were affixed to the custom mounting bracket such that images were captured in a horizontal orientation. Exposure was set to 1/100 s, aperture set to F8 and ISO set to 400. Each camera was manually focussed on the first plant imaged each day and remained fixed for the remaining plants. Images were captured in JPEG format at 24.2-megapixel resolution and saturation boosted in-camera. Each camera was powered by an AC adaptor (EP-5A; Nikon). The cameras were connected via USB cables to a Windows computer running the open source digiCamControl software (Version 2.1.2; Istvan, 2014) for live offload of captured images into the structured folders required for downstream data processing. Images were also backed up onto SD cards installed in each camera. 120 images were captured of each plant (40 with each camera). 120 images were chosen after initial testing (data not shown) revealed this to provide the best balance between reconstruction quality and reconstruction processing time.

### Semi-automated image processing and 3D reconstruction

Image processing and 3D model reconstruction was conducted using open-source software on a Windows PC (as summarised in Figure 3). A dense 3D point cloud was first generated from captured images using VisualSFM (Version 0.5.26 CUDA; Wu, 2011) and CMVS+PMVS2 (Furukawa and Ponce, 2010) using a modified method of Burgess et al. (2015). Processing parameters were adjusted (in the nv.ini configuration file of the VisualSFM working folder; provided as a supplementary file) from the default settings to optimise the reconstruction of chickpea plants (i.e. to provide more detail of the small leaves and thin branches that were not reconstructed in the point clouds using the configuration for wheat of Burgess et al., 2015). Briefly, for CMVS *max_images* was set to 120, for PMVS *min_images* was set to 4, *csize* set to 1, *threshold* set to 0.45 and *wsize* set to 12. Point cloud generation was automated using a Windows batch file (provided as a supplementary file).

**Figure 3.**
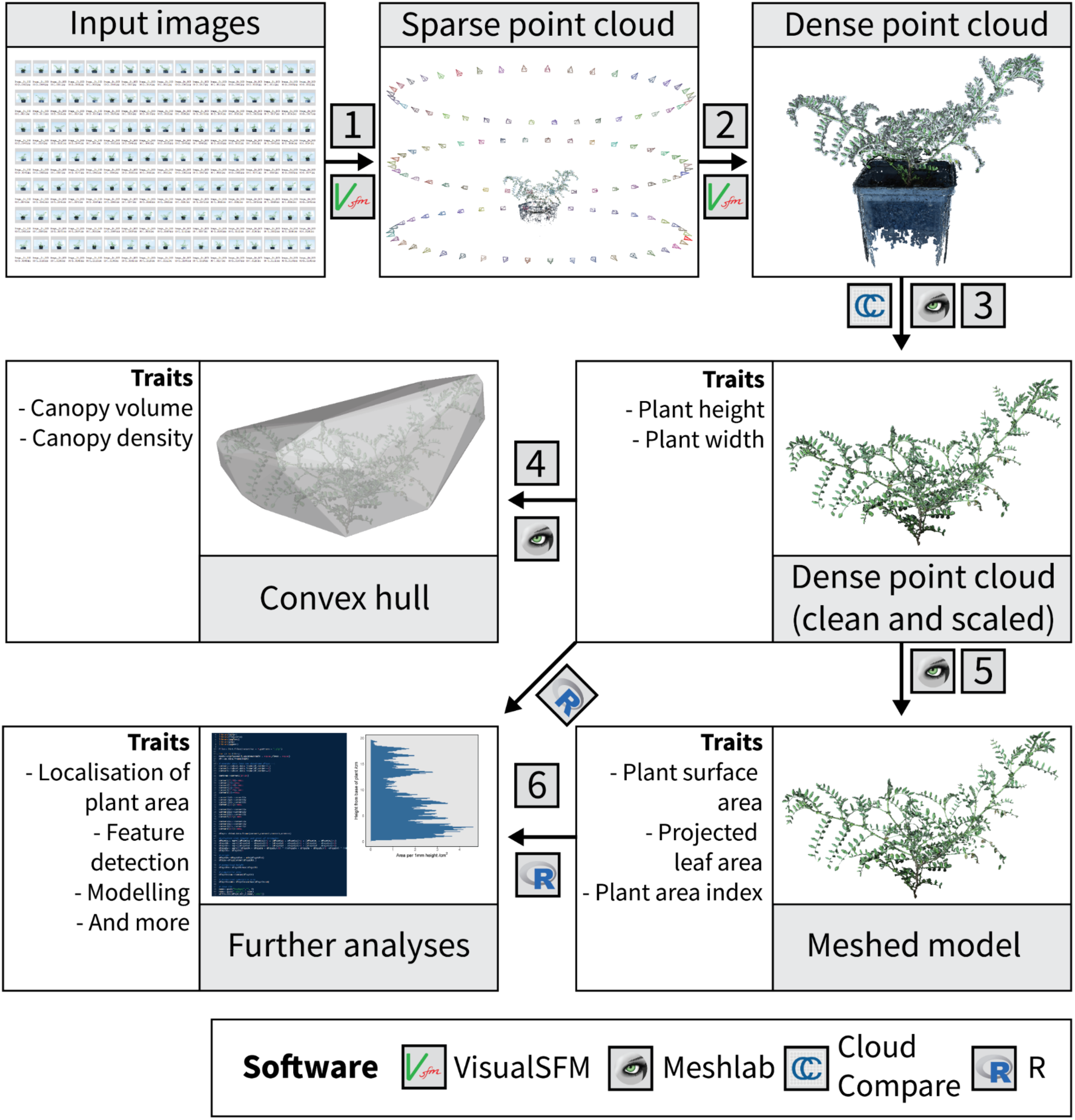
Open source data processing pipeline. **(1)** Captured images are used to generate a sparse point cloud, **(2)** which is then used to generate a dense point cloud. **(3)** Dense point clouds are manually cleaned and scaled, and then used to generate either **(4)** a convex hull or **(5)** a meshed model. **(6)** The scaled point cloud and the meshed model can then be used for further analyses. Note that all but step (3) can be automated in Windows using batch files or, in the case of (6), using R scripts.

Dense point clouds were scaled (using the width of the pot as a reference), denoised based on colour (removing all but the green/brown points), reoriented (such that the ground was parallel to the X-Y plane) and any remaining non-plant points removed manually in Meshlab (Version 2020.06; Cignoni et al., 2008). Statistically outlying points were then removed using the statistical outlier removal (SOR) feature of CloudCompare (Version 2.11.0; GPL software). The remaining points were sub-sampled using Poisson disk sampling (Explicit Radius = 0.5, Montecarlo oversampling = 20; Corsini et al., 2012). A meshed model was created from the sub-sampled point cloud using a ball pivoting algorithm (default settings; Bernardini et al., 1999) and any large holes in the meshed model filled with the close holes feature (max size to be closed = 50). All but the scaling, manual removal of non-plant points and outlier removal were run in a consistent and automated fashion using Meshlab scripts and a Windows batch file (provided as supplementary files). Meshed models consisted of *n* triangles with 3D coordinates of the *i*th triangle given by a vector (*x*_i1_, *y*_i1_, *z*_i1_, *x*_i2_, *y*_i2_, *z*_i2_, *x*_i3_, *y*_i3_, *z*_i3_), where *x* and *y* correspond to coordinates parallel to the ground and *z* corresponds to height above the ground.

Analyses of geometric features (height, max width, etc.) and plant surface area were performed using the base functions in Meshlab. The surface area from Meshlab was divided by 2 to provide a “one-sided” area, which is referred to herein as total surface area. Canopy volume was measured in Meshlab after fitting a convex hull to meshed model. A top down orphographic projection of the model was exported as an image file and processed in ImageJ (Fiji 1.52p; Schindelin et al., 2012) to estimate projected plant area. Plant area index (PAI) was calculated as total surface area/projected plant area. Week-to-week relative growth rates (RGR) for total surface area were derived for each plant as per Pérez-Harguindeguy (2013), using equation 1:

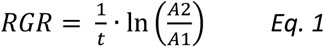

Where *t* is the time between measurement of leaf areas *A*1 and *A*2.

An R script was written to calculate the area of each individual triangle making up the surface of the meshed model and then to calculate plant surface area as a function of height. The script uses the *png* (version 0.1.7; Urbanek, 2013), *rgl* (version 0.100.54; Adler & Murdoch, 2020), *Rvcg* (version 0.19.1; Schlager, 2017) and *tidyverse* (version 1.3.0; Wickham et al., 2019) R packages. Briefly, the length of the i^th^ triangle’s edges (*A*_i_, *B*_i_ and *C*_i_) is first calculated using the XYZ coordinates of its three vertices {*x*_i1_, *y*_i1_, *z*_i1_; *x*_i2_, *y*_i2_, *z*_i2_; and *x*_i3_, *y*_i3_, *z*_i3_}, using equations 2 – 4:

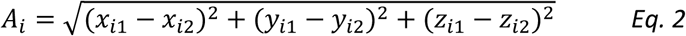

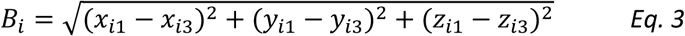

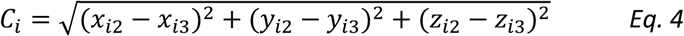

The area of the i^th^ triangle (*S*_i_) is then calculated using lengths *A*_i_, *B*_i_ and *C*_i_ using equation 5:

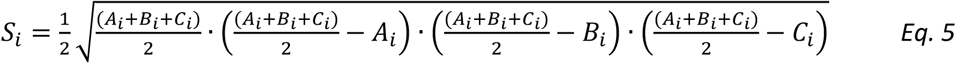

The script outputs a visual summary of plant surface area as a function of height, as well as a comprehensive .CSV file that contains extracted parameters (XYZ coordinates for the vertex of each triangle, XYZ coordinates of the centre of each triangle, the area of each triangle, etc.) from each reconstruction. This R script is provided as a supplementary file. For comparisons across genotypes, for each plant, height data was normalised based on the overall plant height and area data was normalised based on total surface area.

### Validation measurements

The height of each plant was measured using a ruler, from the base of the stem to the highest point of the canopy. Plants were then destructively harvested, the harvested plant material laid flat on a large sheet of white paper and an image taken from above using a DSLR camera (Canon EOS R; Canon Inc., Tokyo, Japan) mounted to a tripod for validation of total surface area (representative images used for ground truthing are presented in Figure S3). A ruler was included in the image for scaling. Lens corrections were first performed on the captured images in Adobe Photoshop (Adobe Inc., San Jose, CA, USA) to remove distortion and then images were analysed using ImageJ (Fiji 1.52p; Schindelin et al., 2012) to obtain measurements of total plant surface area.

### Statistical analyses

Statistical analyses were performed in R (R Core Team, 2019). For validation data, linear regressions models were plotted to visually compare conventional and 3D scanner measurements. Root mean squared error (RMSE) and mean absolute percentage error (MAPE) were calculated using base R and the MLmetrics package (version 1.1.1; Yan, 2016) respectively. Spearman rank correlation coefficient (ρ) was used to statistically analyse the regressions. Analysis of variance (ANOVA) was used to determine whether regression models differed statistically across genotypes. For representative growth data, statistical comparisons across genotypes were analysed using a repeated measures ANOVA with post-hoc Tukey’s HSD test using the *emmeans* package in R (version 1.4.7; Lenth et al., 2020). Normalised area distribution data were analysed statistically using a non-parametric ANCOVA using the *sm* package (version 2.2-5.6; Bowman and Azzalini, 2018). All regressions and representative data were visualised using *ggplot2* in R (version 3.3.2; Wickham, 2016).

## Results

### Reconstruction validation

The 3D reconstructions provided very reliable estimates of plant height and total surface area (Figure 4), both with an R^2^ > 0.99 and Spearman rank correlation coefficient (ρ) > 0.99 when compared to validation measurements. Height was slightly underestimated, with measurements from 3D reconstructions approximately 4% lower than validation measurements, yet there was little variation in this relationship (R^2^ = 0.999, RMSE = 5.45 mm, MAPE = 4.4%, ρ = 0.992, *p* < 0.001) and it was consistent across all studied genotypes (*p* > 0.05; Table S2). Plant surface area measurements were estimated within 0.5% (R^2^ = 0.990, RMSE = 26.85 cm^2^, MAPE = 9.1%, ρ = 0.992, *p* < 0.001), although there was more overall variation in estimates and the validation relationship varied slightly across genotypes (*p* < 0.05; Table S2). Specifically, the surface area of the breeding lines grown outdoors was slightly over-estimated when compared to ground truthing measurements. This was likely caused by smaller, more curled up leaves that were not correctly assessed by ground truthing measurements, which assume all leaves are laid on a two-dimensional plane (for an example see Figure S4). The MAPE in surface area estimates for commercial cultivars (excluding breeding lines) was 7.2%, whilst for the breeding lines the MAPE was 12.3%.

**Figure 4.**
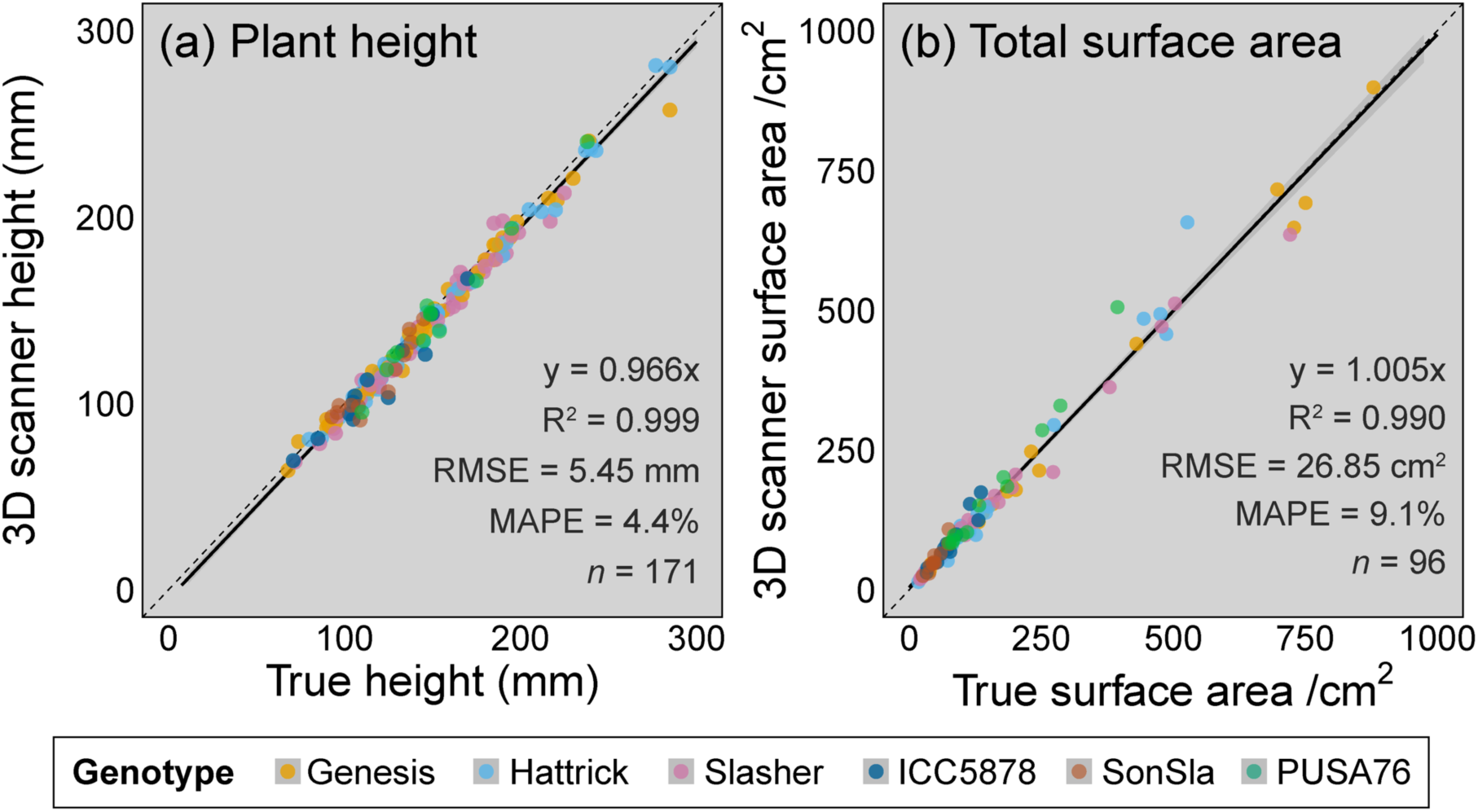
Validation of **(a)** plant height and **(b)** total surface area against conventional measurements. The black lines are linear regressions, the grey shaded region is the standard error. Each point represents an individual plant. Colours represent the seven different genotypes used for validation: Genesis Kalkee (yellow), PBA Hattrick (light blue), PBA Slasher (pink), ICC5878 (dark blue), SonSla (orange) and PUSA76 (green). Please see Table S2 for details of genotype-specific regression models.

### Representative growth data

The 3D scanner allowed us to accurately assess a variety of canopy traits as the plants grew (Figure 5). Whilst there was some variation across individual plants and chickpea genotypes, general trends in growth were clear and easily recovered from 3D reconstructions. Height increased rapidly to a median of 101.2 mm in the first week after germination and then increased more gradually to 191.1 mm five weeks post-germination (Figure 5a). Projected plant area, total surface area and canopy volume all showed characteristic exponential growth curves (Figures 5b, 5c and 5d). Projected plant area increased from a median of 17.0 cm^2^ one week after germination to a median of 220.9 cm^2^ five weeks post-germination, total surface area rose from 37.8 cm^2^ to 415.9 cm^2^ in the same period, and canopy volume from 233 cm^3^ to 14575 cm^3^. Plant area index was not found to vary greatly during the growth of the plants, with a median of 1.91 m^2^ m^−2^ one week after germination and a median of 1.87 m^2^ m^−2^ five weeks after germination. Week-to-week RGR were greatest between weeks 1 and 2, with leaf area increasing on average 84.1% ± 4.4% during this period, dropping to 56.9% ± 4.4%, 51.4% ± 4.0% and 64.1% ± 7.0% between weeks 2 and 3, weeks 3 and 4, and weeks 4 and 5 respectively. Whilst there was some variation in these growth-related traits across individual plants, we found no statistically significant differences across genotypes (*p* > 0.05). Overall variation increased as the plants grew, with some apparent divergence across genotypes in the latter weeks of the experimental period. For example, standard error represented only 10.6% of the mean total surface area in week one whilst it represented 15.5% in week five, with similar trends for the other traits.

**Figure 5.**
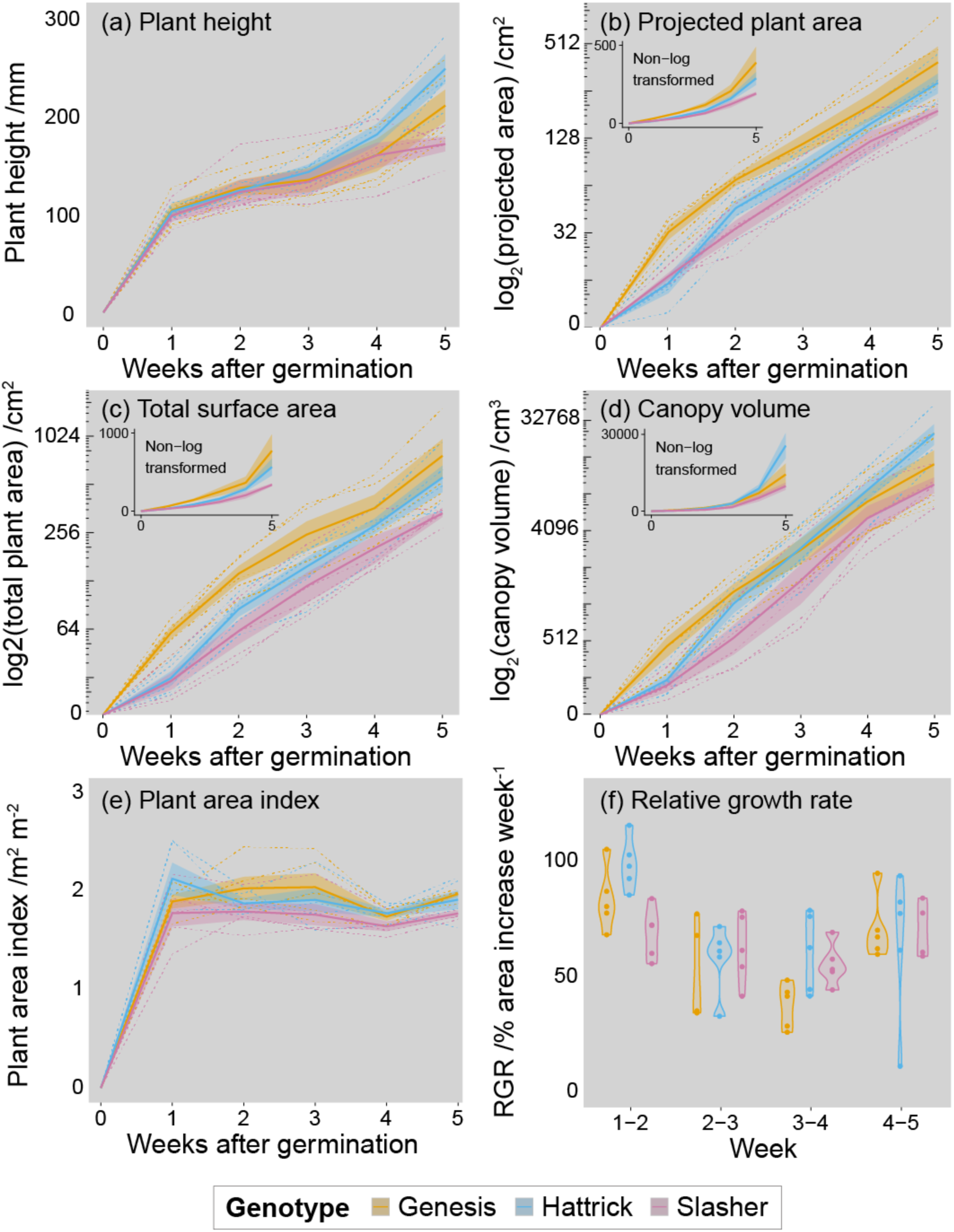
Representative growth data from the 3D scanner, showing **(a)** plant height, **(b)** projected plant area, **(c)** total plant surface area, **(d)** convex hull canopy volume, **(e)** plant area index and **(f)** week-to-week area based relative growth rate (RGR) across five weeks. In panels a-e, solid lines and shaded regions represent genotype means ± SE (n = 15); whilst dashed lines represent individual plants. The main graphs of (b), (c) and (d) are log transformed, with non-log transformed data shown in the inset graphs. In (f) the violins represent the range of RGR for each genotype each week, points are individual plants. Colours represent the three commercial genotypes Genesis Kalkee (yellow), PBA Hattrick (blue) and PBA Slasher (pink).

### Vertical distribution of plant surface area

Further analyses in R enabled us to retrieve detailed data about the distribution of plant surface area as a function of plant height in an automated and repeatable fashion. The visual summaries presented in each panel of Figure 6 are directly outputted from R. These visual summaries provide a fast and semi-quantitative method of assessing how individual plants are partitioning surface area (and by proxy, their biomass). For example, in the representative data shown in Figure 6, the Genesis Kalkee, PBA Hattrick, ICC5878 and PUSA76 plants (Figures 6a, 6b, 6d and 6f respectively) assign most of their plant area to the lower canopy; the PBA Slasher plant (Figure 6c) has a relatively sparse canopy and the SonSla plant (Figure 6e), albeit much smaller than the others, appears to have two discrete canopy layers.

**Figure 6.**
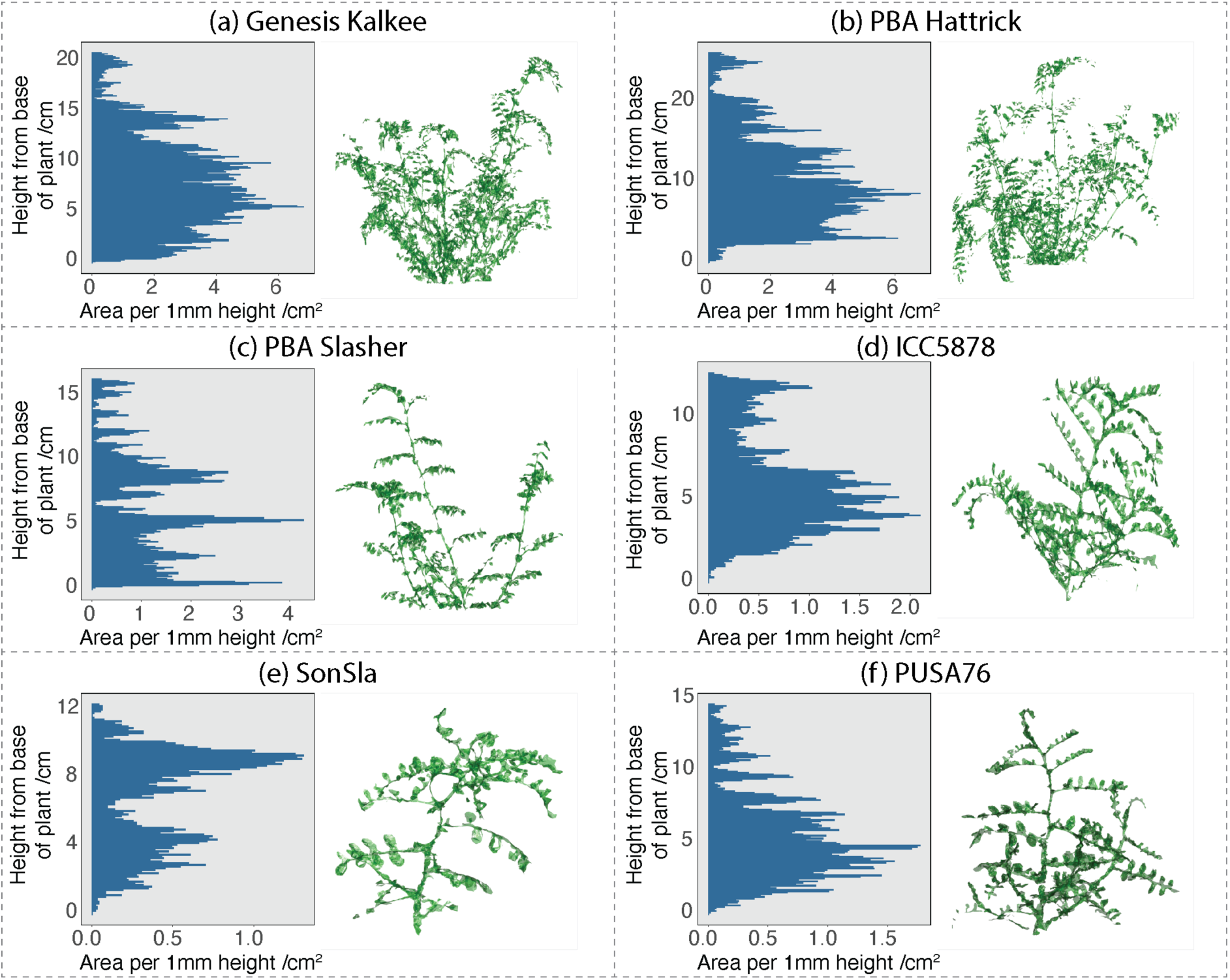
Representative plant surface area distributions for selected individual plants of each genotype (panels **a – e**) at five weeks post-germination. In each panel is (**left**) a graphical summary of the leaf area per mm of height, where each bar represents the sum area of all mesh triangles with centres lying inside each 1 mm z-axis cross section, and (**right**) a 2D visual representation of the meshed model of the plant. Note that the scales of both the graph and the models differ across panels a-e due to variation in the size of individual plants.

To make statistical comparisons of relative area distribution data across genotypes, individual plant data was normalised by plant height and total surface area (Figure 7). Genotypes differed significantly in their relative vertical distribution of leaf area (*p* < 0.001), with particularly clear differences found between the breeding lines and commercial cultivars. The commercial cultivars were much denser in the lower half of the canopy, whilst the breeding lines, and in particular line SonSla, were denser in the mid-to upper-canopy.

**Figure 7.**
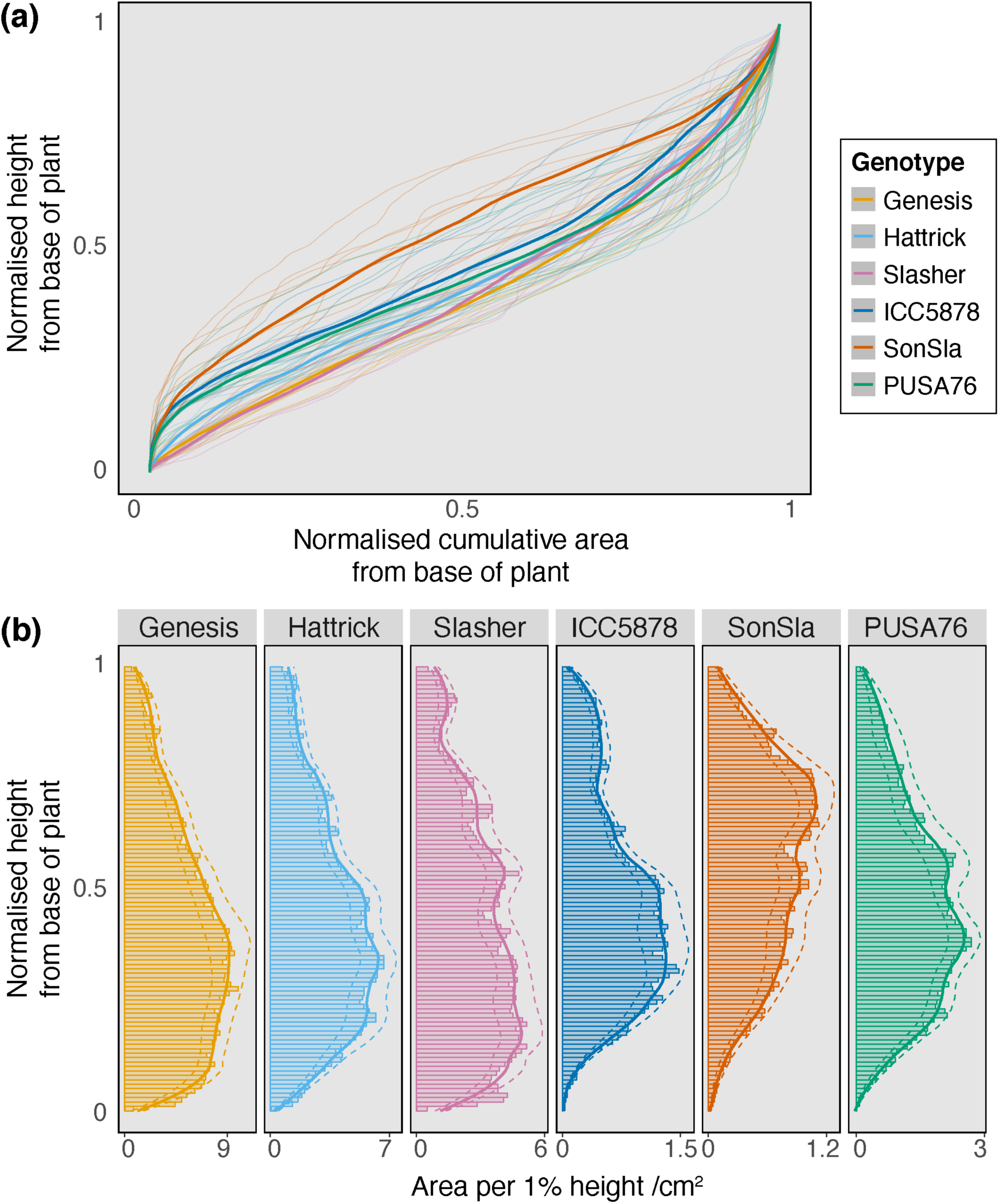
Representative data showing how comparisons of leaf area density distributions could be made across chickpea genotypes by normalising area and height data. All plants presented were imaged at five weeks post-germination. **(a)** is the normalised cumulative surface area from the base to the top of the plant plotted against normalised height. Thick lines represent genotype means (n = 10), thin lines represent individual plants. **(b)** shows surface area versus normalised height, with each bar representing the sum area of all mesh triangles with centres lying inside a 1% z-axis cross section. Values shown by bars are genotype means (n=10) whilst solid and dashed lines represent the genotype mean ± SE respectively fitted with a LOESS function in R.

## Discussion

We have successfully built and validated a low-cost, open source 3D scanner and data processing pipeline to phenotype the architecture and growth trends of individual chickpea plants. Chickpea has leaves that are considerably smaller than most other species that have been phenotyped previously using photogrammetry. In our initial attempts to use the 3D reconstruction workflow developed for wheat by Burgess et al. (2015), we found that there was not enough detail in the 3D reconstructions for accurate measurement of phenotypic traits. However, by modifying key parameters in the reconstruction workflow, we were able to produce reconstructions that provided consistent high-quality data. Validations of height and area measurements from reconstructions against ground truthing measurements highlight the reliability of the system (height, R^2^ > 0.99, MAPE = 4.4%; area, R^2^ = 0.99, MAPE = 9.1%). The accuracy of leaf area estimates is comparable to other photogrammetric estimates reported in the literature for larger leaved plant species (*Brassica napus*, R^2^ = 0.98, MAPE =3.7%, Xiong et al., 2017; maize, sunflower and sugar beet, R^2^ = 0.99, MAPE = 3.9%, Martinez-Guanter et al., 2019; selected houseplant species, R^2^ = 0.99, MAPE = 4.1%, Itakura and Hosoi, 2018; tomato, R^2^ = 0.99, MAPE = 2.3%, Rose et al., 2015). We noted a difference in validation accuracy for plant surface area across genotypes, however, we assigned this to 2D ground truthing measurements underestimating the area of curled up leaves of the outdoor-grown breeding lines, rather than an overestimation of surface area from 3D reconstructions. This underestimation would also explain the greater overall MAPE for area estimates in our study versus other previously studied crops. A similar discrepancy was reported by Bernotas et al. (2019) for *Arabidopsis thaliana*, where top down 2D images consistently underestimated rosette area relative to 3D models that accounted for leaf curvature. In this sense, our 3D reconstructions provide a better estimate of plant surface area than conventional, labour intensive and destructive measurement techniques of chickpea plants.

The results we present here show that photogrammetry could be used as an effective tool to phenotype diversity in plant architecture and growth-related traits across chickpea lines and help to identify novel plant breeding targets. Although we did not find statistically significant differences in architecture traits or growth trends across the three commercial genotypes included in our study, we feel that screening more diverse chickpea lines and continuing to monitor growth for a longer period of time would help to elucidate trends across genotypes. The narrow genetic base of chickpea has hindered improvements in breeding programs in recent years (Thudi et al., 2016). Together with next generation sequencing technologies, the development of new populations selected specifically for the phenotypic investigation of traits of interest could help to address this (Furbank et al., 2020). Even more diversity might be found if we were to investigate traits in wild relatives of cultivated chickpea (Coyne et al., 2020). As the main aim of this study was to evaluate whether photogrammetry could be used to phenotype chickpea, we only monitored the growth of the plants for five weeks post-germination. We did notice there was more variation in architectural traits, both across and within genotypes, as the plants grew larger and future work should seek to assess these traits to plant maturity. The ability to comprehensively assess growth rates of plants to maturity would provide an opportunity to phenotypically screen for highly desirable developmental traits, such as determinacy.

Unlike monocot grain crops such as wheat and barley, chickpea does not have discreet canopy layers, with fruits developing across the whole plant. As such, the optimum light environment for productivity of chickpea canopies will be quite different to that of wheat. The indeterminate nature of chickpea likely shifts this optimum further still, as leaves lower in the canopy will remain photosynthetically active for longer. Modelling could allow us to determine the theoretical optimum light environment and then by running ray tracing simulations with our 3D reconstructions we could determine how close current chickpea architecture is to this optimum. A number of recent studies have used similar approaches to simulate the canopy light environment of other crop species, often coupling this to a photosynthetic model to estimate potential plant productivity (intercropped millet and groundnut, Burgess et al., 2017; sugarcane, Wang et al., 2017; and wheat, Townsend et al., 2018). All three of these modelling studies were conducted using the fastTracer software developed for rice (Song et al., 2013), which we were advised would be very computationally demanding and time consuming to run with our chickpea models due to their complexity (A. J. Burgess, personal communication). There are however some new GPU-based ray tracing software packages that seem promising for simulating the light environment of architecturally complex crops such as chickpea, including the comprehensive and scalable Helios framework developed by Bailey (2019) and the full spectral ray tracer of Henke and Buck-Sorlin (2017). Technological advances such as these will help drive forward our understanding of spatiotemporal variation in the chickpea canopy light environment.

The method we present here provides very reliable estimates of overall plant surface area and other plant traits from whole chickpea plants. We were able to dissect each reconstruction into its component mesh triangles and investigate how plant surface area is distributed relative to plant height. However, what we have so far been unable to do is systematically distinguish between leaf, stem or other plant tissue types in the reconstructions. Segmentation of the models in this way would allow us to retrieve more detailed phenotypic information, including the ability to assess partitioning of biomass across plant tissues, accurately assess other phenotypic traits (such as leaf angles and leaf numbers) and even aid in yield prediction (Shi et al., 2019). Automatic segmentation of 3D models has been achieved in other plant species with larger leaves using several approaches. Itakura & Hosoi (2020) were able to segment individual leaves of a number of broad-leaved houseplant species using a combined attribute-expanding and simple projection segmentation technique. While they retrieved very accurate estimates of leaf area (R^2^ = 0.99, MAPE = 4.1%) using this method, we feel that it would be highly unlikely to work with comparatively tiny chickpea leaves. Another approach would be to use a machine learning algorithm to segment different plant tissues based on pre-trained models. Ziamtsov & Navlakha (2019) recently developed an open-source software package called P3D for this explicit purpose. In their work, they showed P3D to segment leaves and stems in point clouds of tomato and tobacco with > 97 % accuracy. We attempted to use P3D to segment our chickpea models with limited success (data not shown), although this was likely due to the use of the default P3D training datasets developed with larger leaved species. We hope that in the future, with more relevant annotated training datasets, this segmentation technique could also work for chickpea. We provide the full complement of our processed point clouds and meshed models to aid in the development of these training datasets.

The data processing pipeline we have presented here, whilst all open source, does rely on a relatively powerful computer. Specifically, reliable reconstruction of a dense point cloud using PMVS takes a very long time if computer resources (CPU processing power and memory) are limiting. The smaller leaves of chickpea necessitated higher resolution photogrammetry than was needed for the reconstructions of wheat by Burgess et al. (2015). For our reconstructions on a desktop computer with a 16 core/32 thread 3.5 GHz CPU (Ryzen Threadripper 2950X; AMD Inc., Santa Clara, CA, USA) with 128 Gb 3200 MHz RAM (HyperX Fury; Kingston Technology Corp., Fountain Valley, CA, USA), the generation of a dense point cloud took roughly two hours per plant. We also found that running the reconstruction process from image data stored on a solid-state drive was considerably faster than running from images stored on a traditional hard disc drive. In the past, such computing resources would have been prohibitively expensive for most researchers however this is no longer the case, largely thanks to advances driven by computer gaming technology. Multicore computing is now the norm, even in portable laptop computers, and high capacity memory and fast solid-state storage are now reasonably priced.

On the topic of cost, our imaging set up cost roughly AU $1300, considerably less than commercially available alternatives that offer similar data quality. Panjvani et al. (2019) recently developed a comparably priced (US $400) DIY LIDAR system for 3D scanning of individual plants, however the quality of leaf area estimates was considerably less than ours (R^2^ < 0.6 against ground truthing data, MAPE = 31.5%). By far the most expensive part of our set up was the cameras. In our method presented here, we used three DSLR cameras however we must highlight that the method can also be adapted to work with just one camera, substantially reducing cost. In our early testing, we used just one camera and rotated the plant three times, with the camera manually repositioned from one mounting point of the camera bracket to the next between each rotation. Whilst this took us longer to capture the image sets, we did not notice any reduction in data quality. It may also be possible to use cheaper cameras. Martinez-Guantar et al. (2019) used a regular point and shoot camera for the 3D reconstruction of maize, sunflower and sugar beet plants, with an R^2^ > 0.99 for both height and leaf area estimates compared against ground truthing measurements. Paturkar et al. (2020) show that even a mobile phone can be used for image capture, with 3D reconstructions of chilli plants giving an R^2^ > 0.98 for estimates of both height and leaf area. These technological advances and reductions in cost mean that photogrammetric techniques are more accessible than ever before to the plant phenotyping community. The increased availability of these technologies will allow for the adoption of data driven approaches to fundamental and pre-breeding plant science research where this was not possible before.

## Conclusions

Our work has shown that it is possible to use low-cost photogrammetry techniques to accurately phenotype architectural traits and growth habits of individual chickpea plants. We hope that our use of open source software and hardware will allow others to easily reproduce our method and to develop it further. In particular, there is a need to test whether photogrammetry reconstructions of chickpea could be used for simulations of the canopy light environment and whether they could be automatically segmented into different plant organs using deep learning algorithms. There is a need for higher yielding, environmentally friendly and stress tolerant chickpea varieties with increasing demand for high quality pulse protein worldwide. The use of novel phenotyping tools and associated data analytics should assist us in identifying traits of interest and allow us to explore diversity in these traits so that breeders can make informed breeding decisions.

## Data availability

All 3D meshed models and scaled dense point clouds used in this study are openly available on Zenodo (DOI: 10.5281/zenodo.4018242).

## Acknowledgements

We would like to thank Dr Alexandra Burgess and Dr Jonathon Gibbs from The University of Nottingham for their useful advice and initial assistance in running the 3D reconstruction software. We thank Dr Angela Pattison for supplying chickpea seed and Dr Sonam Tashi for assistance growing the plants. This work was supported by the Australian Research Council (ARC), Industrial Transformation Research Hub —Legumes for Sustainable Agriculture (IH140100013) and the Grains Research and Development Corporation.

## Supplementary tables

**Table S1.** Chickpea genotypes used to validate the 3D scanner. These lines were chosen based on their contrasting canopy heights, growth habits and growth rates. Note that some information is lacking for breeding lines.

| Genotype | Type | Plant height | Growth habit | Vigour |
| --- | --- | --- | --- | --- |
| Genesis Kalkee | Kabuli | Medium/tall | Erect | Mid |
| PBA Hattrick | Desi | Tall | Erect | Mid |
| PBA Slasher | Desi | Medium/short | Spreading | Late |
| ICC5878 | Desi | Short | - | - |
| SonSla | Desi | - | Prostrate | - |
| PUSA76 | Desi | - | - | Early |

**Table S2.** Genotype specific regressions for validation measurement. Significance values refer to an ANOVA run to determine whether there were differences across genotypes.

| Genotype | Regression | R<sup>2</sup> | n |
| --- | --- | --- | --- |
| **Height validation<sup>n.s.</sup>** |  |  |  |
| Genesis Kalkee | y = 0.993x | 0.999 | 45 |
| PBA Hattrick | y = 0.970x | 0.999 | 45 |
| PBA Slasher | y = 0.963x | 0.999 | 45 |
| ICC5878 | y = 0.941x | 0.996 | 12 |
| SonSla | y = 0.951x | 0.996 | 12 |
| PUSA76 | y = 0.975x | 0.998 | 12 |
| **Area validation\*** |  |  |  |
| Genesis Kalkee | y = 0.986x | 0.997 | 20 |
| PBA Hattrick | y = 1.072x | 0.988 | 20 |
| PBA Slasher | y = 0.941x | 0.994 | 20 |
| ICC5878 | y = 1.122x | 0.980 | 12 |
| SonSla | y = 1.137x | 0.979 | 12 |
| PUSA76 | y = 1.167x | 0.993 | 12 |
| **Significance** |  |  |  |
| <sup>n.s.</sup> = P > 0.05; * = P < 0.05; ** = P < 0.01; *** = P < 0.001. |  |  |  |

## Supplementary figures

**Figure S1.**
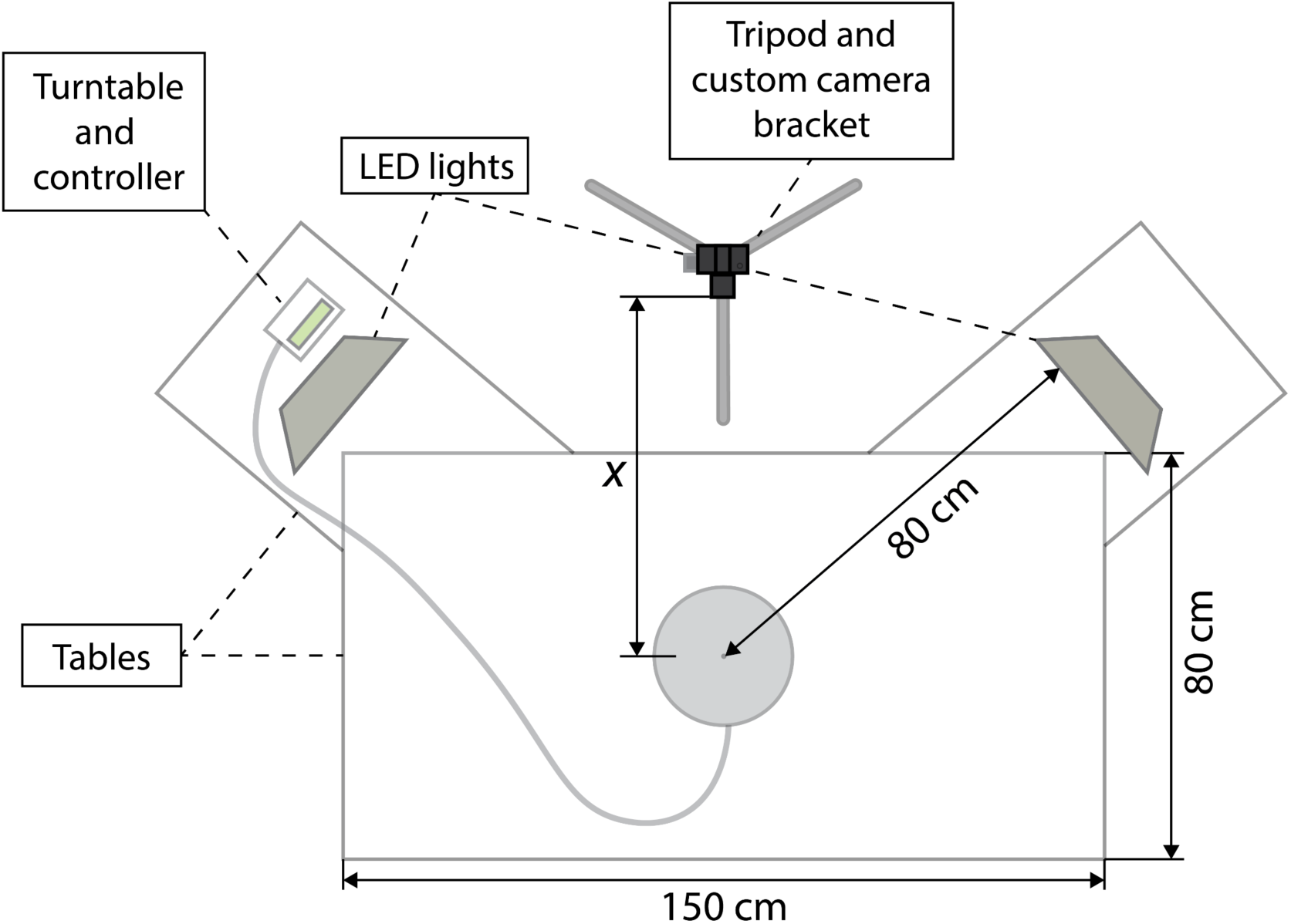
Top down diagram of the laboratory imaging set up. The tripod was moved back from the table (distance *x*) as the plants grew.

**Figure S2.**
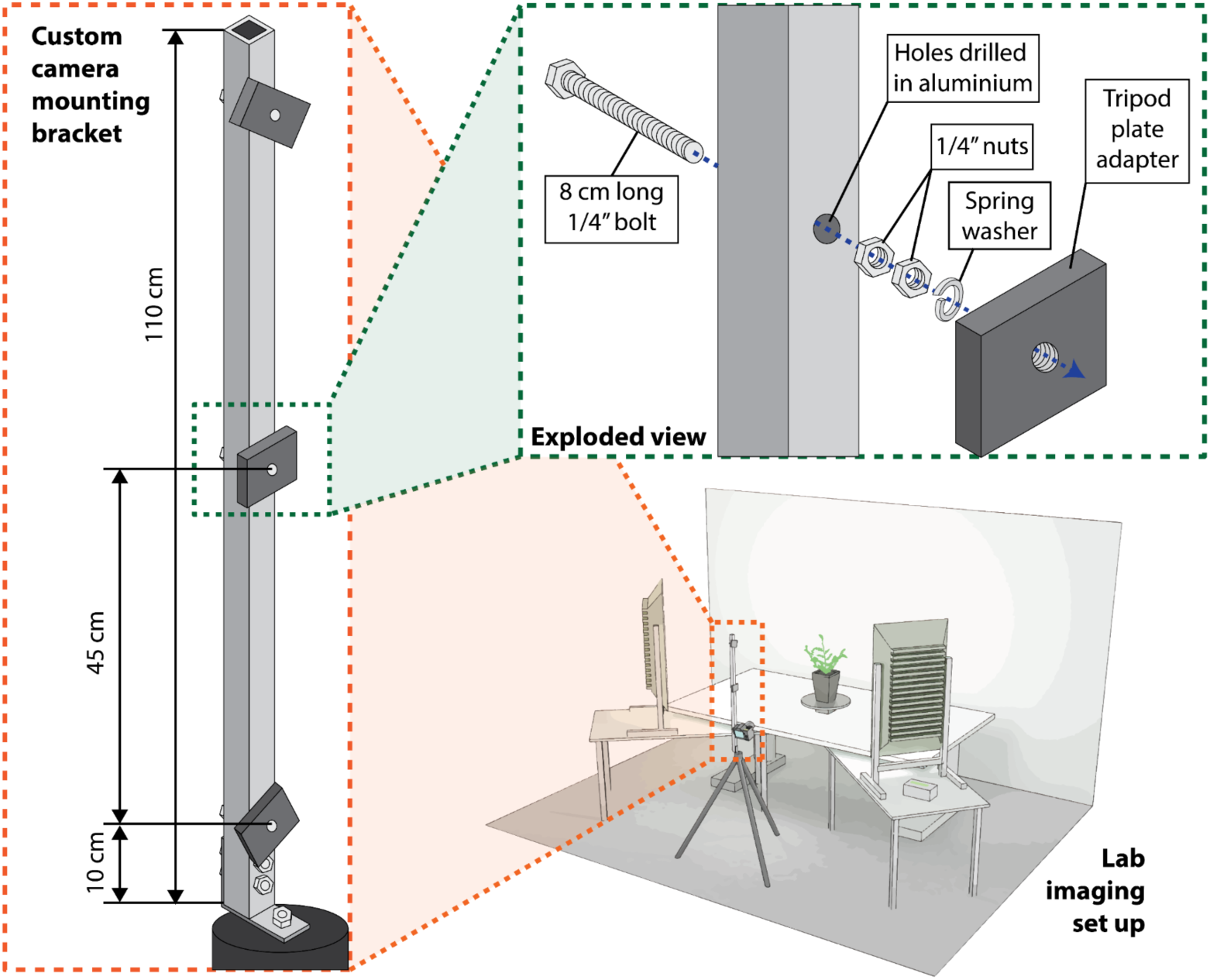
Schematic diagram of the custom camera mounting bracket.

**Figure S3.**
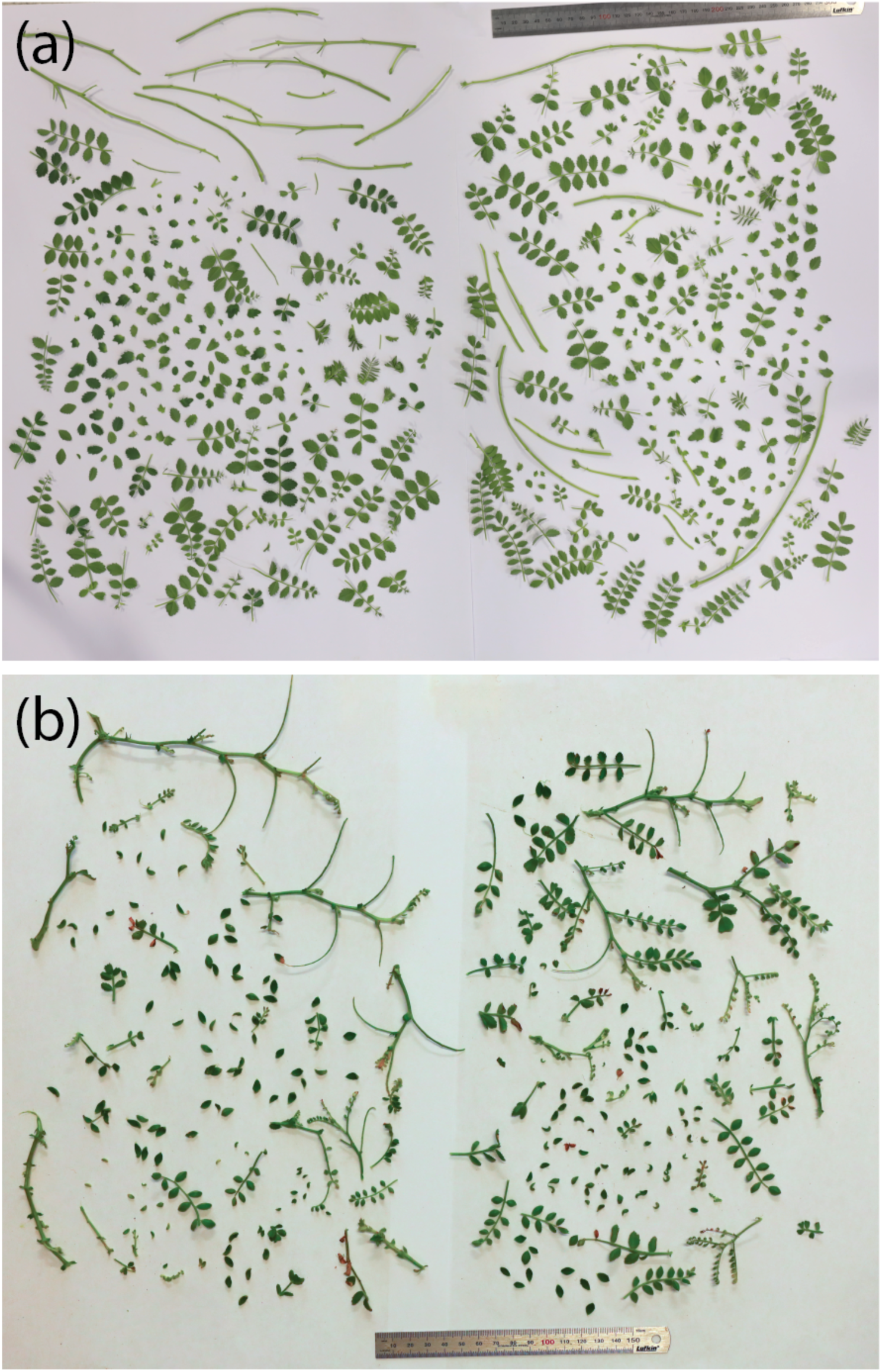
Two representative images of a harvested chickpea plants used for the ground truthing measurement of leaf area. **(a)** is the commercial chickpea cultivar Genesis Kalkee and **(b)** is the breeding line PUSA76.

**Figure S4.**
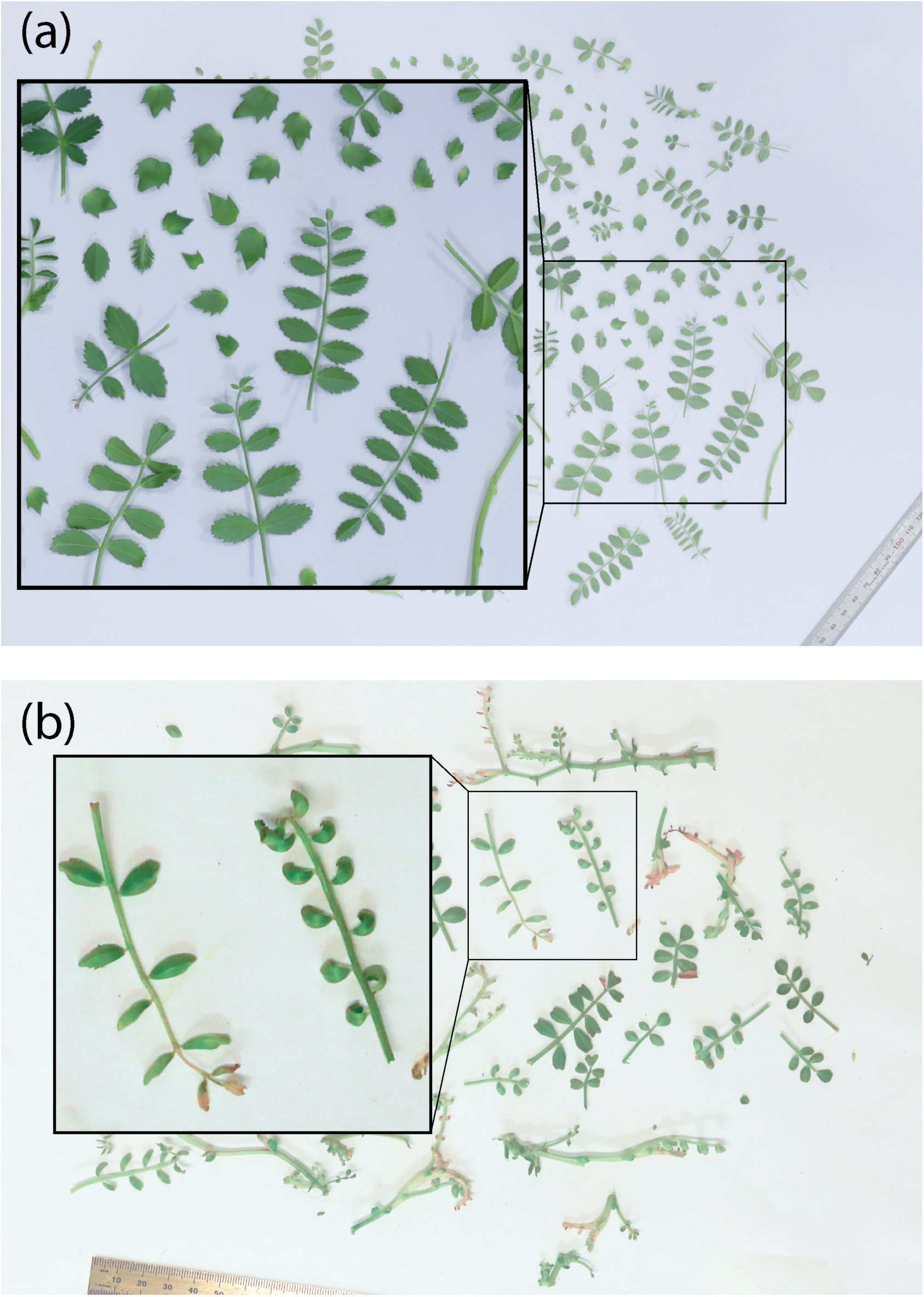
An example of the curled leaves that caused an underestimation of leaf area in ground truthing measurements of the outdoor-grown chickpea breeding lines. **(a)** shows a representative plant of the commercial cultivar, the inset figure shows that most leaves were flat. **(b)** shows a representative plant of breeding line SonSla, the inset figure shows that many leaves were curled or folded to a certain degree.

## List of supplementary processing files

1. *3Dscannerprogram*.*ino* – Arduino program for 3D scanner. Please refer to comments embedded in code to customise capture sequence. Upload to Arduino board using the Arduino IDE software available from arduino.cc.
2. *nv*.*ini* – VisualSFM configuration file for chickpea plants. To use, please replace nv.ini file in the VisualSFM working folder. Please refer to comments embedded in code to customise reconstruction process; this file has optimised parameters for chickpea plants.
3. *VisualSFMbatch*.*bat* – Windows batch file for generation of dense point clouds from image sets. Please ensure you change the filepaths in the code to the folder containing image sequences and the output file path.
4. *Autocleanscript*.*mlx –* Meshlab script to reorient point cloud, remove non-green, non-plant points and remove outlying points.
5. *Meshlabbatchautoclean*.*bat* – Windows batch file to run the “Autocleanscript.mlx” meshlab script on multiple models sequentially. Please ensure you change the filepaths in the code to the folder containing raw dense point clouds .ply files from VisualSFM, the output file path and the file location of the “Autocleanscript.mlx” file.
6. *Meshscript*.*mlx* – Meshlab script to mesh clean and scaled point clouds using a ball pivoting algorithm and close any holes in the resultant meshed model.
7. *Meshlabbatchmesh*.*bat* – Windows batch file to run the “Meshscript.mlx” meshlab script on multiple models sequentially. Please ensure you change the filepaths in the code to the folder containing clean and scaled point cloud .ply files, the output file path and the file location of the “Meshscript.mlx” file. Can also output a text file summary of the model containing dimensions and surface area of the meshed model.
8. *Convexhullscript*.*mlx* – Meshlab script to fit a convex hull to clean and scaled point clouds.
9. *Meshlabbatchconvexhull*.*bat* - Windows batch file to run the “Convehullscript.mlx” meshlab script on multiple models sequentially. Please ensure you change the filepaths in the code to the folder containing clean and scaled point cloud .ply files, the output file path and the file location of the “Convexhullscript.mlx” file. Can also output a text file summary of the model containing convex hull volume.
10. *Plantmodelbreakdown*.*r* – R script for processing leaf area distribution by height. Please refer to comments embedded in code to customise outputs etc.

## Notes

https://doi.org/10.5281/zenodo.4018242

### Competing Interest Statement

The authors have declared no competing interest.

