## Supplementary material for "Open source 3D phenotyping of chickpea plant architecture across plant development": 3Dscannerprogram.ino

```
/* Chickpea 3D scanner

Controls
SELECT: Start/Stop automatic scan.
DOWN: Trigger camera shutter and advance one unit clockwise.
RIGHT: Step one unit clockwise without triggering camera.
LEFT: Step one unit counter clockwise without triggering camera.
UP: Cycle through options for number of exposures per revolution.

*/

/*Load libraries*/
#include <LiquidCrystal.h>
#include <DFR_Key.h>
#include "DRV8825.h"


/*Input stepper motor properties and load driver*/
#define MOTOR_STEPS 200
#define RPM 1
#define MICROSTEPS 32
#define DIR 12
#define STEP 13
#define MODE0 0
#define MODE1 1
#define MODE2 2
DRV8825 stepper(MOTOR_STEPS, DIR, STEP, MODE0, MODE1, MODE2);


/*Load display and keypad driver and define pins in use*/
LiquidCrystal lcd(8, 9, 4, 5, 6, 7); //pin assignments for SainSmart LCD Keypad Shield
DFR_Key keypad; // initialize keypad

/*Input variables for turntable*/
int localKey = 0;
int lastKey = 0;
int keyChange = 1; //set true when key is released.  Prevents double triggers.
int stepChoices [] = {40,60,80}; //different numbers of exposures per revolution which may be selected.  Higher number = smaller angle between successive shots
int stepIndex = 0;  // count exposures starting at 1
int numChoices = 5;  //number of step choices
int runFlag = 0; //is scanner running
int stepCount =1;  // count exposures starting at 1
unsigned long startWait;  //time we start the wait timer
unsigned long currentTime; // current time
long preWait = 7000;  // pre exposure pause in ms.  Allows the specimen to settle before exposure.
long postWait = 500;  // post exposure pause in ms.  Allows time for the exposure to finish before moving.
int waitFlag = 0; // 0=ready to move 1=pre-exposure wait 2=post-exposure wait

/*Define camera shutter pin and shutter speed in ms*/
int Camerapin = 11;
long Cameradelay = 500;

/*Define end buzzer pin and length of buzzer chime in ms*/
int Buzzerpin = 3;
long Buzzerdelay = 2000;

/* Startup program */
void setup() 
{  
  lcd.begin(16, 2);
  lcd.clear();
  lcd.setCursor(0, 0);
  lcd.print("Chickpea 3D Scanner");
  lcd.setCursor(0, 1);
  lcd.print("by W.T.Salter");
  pinMode(Camerapin, OUTPUT);
  digitalWrite(Camerapin, LOW);
  pinMode(Buzzerpin, OUTPUT);
  digitalWrite(Buzzerpin, LOW);
  delay(300);
  lcd.clear();

stepper.begin(RPM, MICROSTEPS);

}

/* Main program */
void loop(){
 
  //
    lcd.setCursor(0,0);
    lcd.print("Steps:");
    lcd.setCursor(6,0);
    lcd.print("   ");
    lcd.setCursor(6,0);
    lcd.print(stepChoices[stepIndex]);
    lcd.setCursor(0,1);
    if (runFlag == 1)
    {
      lcd.print("run    ");
      
    }
    else
    {
      lcd.print("stop    ");
    }
  
  
   localKey = keypad.getKey(); // read keypad
  
  if (localKey != SAMPLE_WAIT)
  {
    
    
    
    if (localKey == 0) // key has been released
      {
       keyChange =1; // the next key data represents a new key press
      }
    
    if (localKey == 1 && keyChange == 1) //select=start sequence
      {
        if (runFlag == 0 && keyChange ==1)
        {
          runFlag = 1;
          waitFlag = 0;
          keyChange = 0;

        }
        if (runFlag == 1 && keyChange ==1)
        {
          runFlag = 0;
          stepCount = 1;  //uncomment to reset stepCount every time the auto scan is stopped.
          keyChange = 0;

        }
      }
      
    if (localKey == 4 && keyChange == 1) //down=manual mode
      {
       keyChange =0; 
       lcd.setCursor(0,1);
       lcd.print("manual");
       digitalWrite(Camerapin, HIGH); //trigger exposure
       delay(Cameradelay);
       digitalWrite(Camerapin, LOW);
 
      }
      
    if (localKey == 3 && keyChange == 1 && runFlag ==0) //up cycle through angle choices
      {
       keyChange =0;
       if (stepIndex < numChoices -1)
         {
           stepIndex ++;
         }
       else
         {
           stepIndex = 0;
         }
      }
      
      
      
    if (localKey == 2 && keyChange == 1) //left step
      {
       keyChange =0;
      lcd.setCursor(0,1);
      lcd.print("CCW step");
      stepper.move(-MOTOR_STEPS*32/stepChoices[stepIndex]);

   
      }
      
    if (localKey == 5 && keyChange == 1) //right step
      {
       keyChange =0;
      lcd.setCursor(0,1);
      lcd.print("CW step");
       stepper.move(MOTOR_STEPS*32/stepChoices[stepIndex]);

    
      }

}


    if (runFlag == 1) //capture sequence is running
    {
     
      
      if (stepCount > stepChoices[stepIndex]) //capture process is complete, sound buzzer then reset counters. 
      {
        runFlag = 0; // stop sequence
        lcd.setCursor(9,1);
        lcd.print("Done   ");
        digitalWrite(Buzzerpin, HIGH);
        delay(Buzzerdelay);
        digitalWrite(Buzzerpin, LOW);
        stepCount = 1;
        
      }


//This interrupt based time delay allows us to still receive keypad input during the delay

      currentTime = millis();

      if (waitFlag == 2) // wait after triggering shutter before moving motor
      {
         if (currentTime - startWait >= postWait) //wait time has expired
         {
           waitFlag = 0; //done waiting
         }
      }
      

      if (waitFlag == 0)  // advance stepper and start wait timer
      {
        startWait = millis();
        waitFlag = 1; // start preshutter wait
        lcd.setCursor(9,1);
        lcd.print("Exp#:   ");
        lcd.setCursor(13,1);
        lcd.print(stepCount);
        stepper.move(MOTOR_STEPS*32/stepChoices[stepIndex]);  //advance stepper
      }

      
      if (waitFlag == 1) // when preshutter wait expires trigger shutter
      {

        if (currentTime - startWait >= preWait) //wait time has expired
        {
        digitalWrite(Camerapin, HIGH); //trigger exposure
        delay(Cameradelay);
        digitalWrite(Camerapin, LOW);
        startWait = millis(); // restart wait timer
        waitFlag = 2; // initiate post shutter wait
        stepCount ++; 
                }
         
    }
   }
}
```
