## Supplementary material for "Open source 3D phenotyping of chickpea plant architecture across plant development": Meshlabbatchautoclean.bat

```
rem update filepath to folder containing Meshlab executable
cd C:\Program Files\VCG\MeshLab

rem update to include folder containg .ply dense point cloud from VisualSFM, output file destination and filename, and folder containing Meshlab .mlx script. Add more if needed, processing will proceed sequentially.
start /wait cmd /c meshlabserver -i C:\Users\USERNAME\Documents\3D_Scanner_Image_Folder\UncleanedPointClouds\Plant1.ply -o C:\Users\USERNAME\Documents\3D_Scanner_Image_Folder\AutocleanedPointClouds\Plant1.ply -s C:\Users\tamsa\Documents\3D_Scanner_Image_Folder\Meshlab_Scripts\Autocleanscript.mlx  
start /wait cmd /c meshlabserver -i C:\Users\USERNAME\Documents\3D_Scanner_Image_Folder\UncleanedPointClouds\Plant2.ply -o C:\Users\USERNAME\Documents\3D_Scanner_Image_Folder\AutocleanedPointClouds\Plant2.ply -s C:\Users\tamsa\Documents\3D_Scanner_Image_Folder\Meshlab_Scripts\Autocleanscript.mlx  
start /wait cmd /c meshlabserver -i C:\Users\USERNAME\Documents\3D_Scanner_Image_Folder\UncleanedPointClouds\Plant3.ply -o C:\Users\USERNAME\Documents\3D_Scanner_Image_Folder\AutocleanedPointClouds\Plant3.ply -s C:\Users\tamsa\Documents\3D_Scanner_Image_Folder\Meshlab_Scripts\Autocleanscript.mlx  
start /wait cmd /c meshlabserver -i C:\Users\USERNAME\Documents\3D_Scanner_Image_Folder\UncleanedPointClouds\Plant4.ply -o C:\Users\USERNAME\Documents\3D_Scanner_Image_Folder\AutocleanedPointClouds\Plant4.ply -s C:\Users\tamsa\Documents\3D_Scanner_Image_Folder\Meshlab_Scripts\Autocleanscript.mlx  
start /wait cmd /c meshlabserver -i C:\Users\USERNAME\Documents\3D_Scanner_Image_Folder\UncleanedPointClouds\Plant5.ply -o C:\Users\USERNAME\Documents\3D_Scanner_Image_Folder\AutocleanedPointClouds\Plant5.ply -s C:\Users\tamsa\Documents\3D_Scanner_Image_Folder\Meshlab_Scripts\Autocleanscript.mlx  
```
