## Supplementary material for "Open source 3D phenotyping of chickpea plant architecture across plant development": Meshlabbatchmesh.bat

```
rem update filepath to folder containing Meshlab executable
cd C:\Program Files\VCG\MeshLab

rem update to include folder containg cleaned and scaled .ply point clouds (note thjat this requires some manual steps), output file destinations and filenames, and folder containing Meshlab .mlx script. Add more if needed, processing will proceed sequentially.
start /wait cmd /c meshlabserver -i C:\Users\USERNAME\Documents\3D_Scanner_Image_Folder\AutocleanedPointClouds\Plant1.ply -o C:\Users\USERNAME\Documents\3D_Scanner_Image_Folder\Meshedmodels\Plant1.ply -s C:\Users\USERNAME\Documents\3D_Scanner_Image_Folder\Meshlab_Scripts\Meshscript.mlx  -l C:\Users\USERNAME\Documents\3D_Scanner_Image_Folder\Meshedmodels\Plant1.txt
start /wait cmd /c meshlabserver -i C:\Users\USERNAME\Documents\3D_Scanner_Image_Folder\AutocleanedPointClouds\Plant2.ply -o C:\Users\USERNAME\Documents\3D_Scanner_Image_Folder\Meshedmodels\Plant2.ply -s C:\Users\USERNAME\Documents\3D_Scanner_Image_Folder\Meshlab_Scripts\Meshscript.mlx  -l C:\Users\USERNAME\Documents\3D_Scanner_Image_Folder\Meshedmodels\Plant2.txt
start /wait cmd /c meshlabserver -i C:\Users\USERNAME\Documents\3D_Scanner_Image_Folder\AutocleanedPointClouds\Plant3.ply -o C:\Users\USERNAME\Documents\3D_Scanner_Image_Folder\Meshedmodels\Plant3.ply -s C:\Users\USERNAME\Documents\3D_Scanner_Image_Folder\Meshlab_Scripts\Meshscript.mlx  -l C:\Users\USERNAME\Documents\3D_Scanner_Image_Folder\Meshedmodels\Plant3.txt
start /wait cmd /c meshlabserver -i C:\Users\USERNAME\Documents\3D_Scanner_Image_Folder\AutocleanedPointClouds\Plant4.ply -o C:\Users\USERNAME\Documents\3D_Scanner_Image_Folder\Meshedmodels\Plant4.ply -s C:\Users\USERNAME\Documents\3D_Scanner_Image_Folder\Meshlab_Scripts\Meshscript.mlx  -l C:\Users\USERNAME\Documents\3D_Scanner_Image_Folder\Meshedmodels\Plant4.txt
start /wait cmd /c meshlabserver -i C:\Users\USERNAME\Documents\3D_Scanner_Image_Folder\AutocleanedPointClouds\Plant5.ply -o C:\Users\USERNAME\Documents\3D_Scanner_Image_Folder\Meshedmodels\Plant5.ply -s C:\Users\USERNAME\Documents\3D_Scanner_Image_Folder\Meshlab_Scripts\Meshscript.mlx  -l C:\Users\USERNAME\Documents\3D_Scanner_Image_Folder\Meshedmodels\Plant5.txt
```
