## Supplementary material for "Open source 3D phenotyping of chickpea plant architecture across plant development": nv.ini

```
############################################
#Default ratio between focal length and max(width, height)
#This parameter is very useful for wide angle images
#Change this parameter before launching VisualSFM
param_default_focal_ratio 1.2
#Change this to 0 if the focal length from EXIF is too bad
param_use_exif_focal_length 0
#Enable radial distortion (1 to enable, 0 to disable)
param_radial_distortion 1

#Switch to preemptive matching mode when the number of images is large than
#param_preemptive_switch and preemptive matching is enabled 
#(+subset for command-line or SfM->Pairwise Matching->Use Preemptive Matching) 
param_preemptive_switch 1000
#Number of features used in preemptive matching (h in the 3DV2013 paper)
param_preemptive_size 100
#Threshold for preemptive matching (t_h in the 3DV2013 paper)
param_preemptive_min 4

#Bundle adjustment configuration
param_bundle_full_step 5
param_bundle_full_iteration 100
param_bundle_part_size 20
param_bundle_part_iteration 5
param_bundle_stop_mse 0.25
param_pba_prefer_gpu 0
param_pba_use_schur 0

#Skip visualization data as much as possible (use 1 to save memory)
param_less_visualization_data 1
#The number of images that allows thumbnail textures and pixel loading.
param_visual_data_budget 500
#Enable full resolution rendering for resolution larger than 4096.
param_enable_huge_texture 0

#The number of threads for image undistortion (0 means one-thread per core)
param_undistortion_thread_num 32
#Parameters for using CMVS/PMVS (by Yasutaka Furukawa) 
param_cmvs_max_images 120
param_pmvs_min_image 4
param_pmvs_level 1
param_pmvs_csize 1
param_pmvs_threshold 0.45
param_pmvs_wsize 12
param_pmvs_max_angle 10
#Thread parameter for PMVS: 0 means one-thread per core.
param_pmvs_max_cpus 32

#GPS variance used for GPS-based 3D transformation (beta)
param_gps_variance 10
param_gps_ignore_exif_altitude 1
#Do not transform the NVM or PLY. Save the transformation to aln instead. 
param_write_transformation_to_aln 0

#Thresholds for triangulating a 3D point
param_triangle_angle_min_new 4
param_triangle_angle_min 2
param_forward_motion_threshold 2
#Threshold for adding a point to existing track and making new tracks
param_projection_threshold 320
param_new_track_threshold 320
param_merge_track_threshold 32
param_retriangulate_threshold 3200
param_retriangulate_step 0.25
#Threshold for removing tracks that have large projection errors
param_track_threshold 8
#Filter unstable points (small triangulation angle)
param_remove_unstable_tracks 1

#Threshold for the number of projections when adding a new camera
param_new_cam_projection_min 20
#Threshold for pose estimation
param_pose_estimate_threshold 64
#Adjust pose estimation threshold according to focal lengths
param_pose_estimate_adaptive 1
#Consider very large focal length as wrong
param_pose_estimate_ratio_max 10
param_p3p_inlier_ratio_min 0.4
param_pose_inlier_ratio_min 0.2

#The minimum inlier matches for a pair to be considered correct
param_pair_inlier_num_min 15
#Distance threshold in meter for pair-wise image matching (0 to disable)
param_pair_gps_threshold 0
#Aggressively include all putative matches in reconstruction
param_aggressive_triangulation 0

#The thresholds for selecting a starting pair 
#Homography threshold : the maximum percent of points on a plane
param_init_h_ratio_max 0.4
#Inlier count threshold : the minimum number of matches
param_init_inlier_min 30
#Triangulation angle threshold: minimum average triangulation angle
param_init_angle_min 2

#Reconstruct one model or multiple?
param_search_multiple_models 1
#Merge two models if the number of 3D point correspondences is large enough
param_model_merge_min_matches 10
#Maximum images shared by two 3D models
param_image_reuse_max 5

#Save a temporary NVM file when <Reconstruct Sparse> finishes.
param_model_auto_saving 0
#Write the combined PLY file in ASCII or BINARY format
param_write_ascii_ply 0
#Write camera indices in PLY file
param_write_cameras_in_ply 1

#Some external applications to run from VisualSFM.
#%s will be replaced by the nvm or ply file title.
param_external_tool1 PoissonRecon --verbose --in %s.ply --out %s.poisson.ply --depth 10
param_external_tool2 my_ply_upload_tool %s.ply
param_external_tool3 my_nvm_process_tool %s.nvm

#Select the tool for feature detection 
#0: CPU (depending on param_use_vlfeat_or_lowe),
#1: GLSL-based SiftGPU, 2 CUDA-based SiftPGU,
#3: customized SiftGPU (depending on param_use_siftgpu_customize)
param_use_siftgpu 3
param_use_siftgpu_customize -cuda 0 -fo -1 -v 1 -tc -1 -nomc -maxd 6000
#Use the original binary from Lowe if 0, and VLFeat binary otherwise
param_use_vlfeat_or_lowe 0
#Options for vlfeat sift (--peak-thresh=3.4 seems to be compatible with Lowe)
param_vlfeat_options --peak-thresh=3.4

#Select the device for feature matching 
#0: cpu, 1: glsl, 2+: cuda if compiled in SiftGPU.dll
param_use_siftmatchgpu 2
param_cpu_siftmatch_num_thread 32

#Feature matching parameters
#Max features used in gpu sift matching
param_gpu_match_fmax 1000000
#Max sift matching distance
param_sift_max_dist 0.7
#Max sift matching distance ratio
param_sift_max_dist_ratio 0.8
#Mutual best match or not
param_siftmatch_mbm 1
#Filter the stationary point matching
param_no_stationary_points 0

#Two-view geometry estimation parameters
#Threshold for estimating F in ransac
param_ransac_f_threshold 2
#Threshold for estimating H in ransac
param_ransac_h_threshold 4
#Maximum iterations in ransac
param_ransac_max_sample 1000

#Disable GPU-based computation when remote desktop is detected
#Change this to 0 if VisualSFM incorrectly detects remote desktop
param_check_remote_desktop 0
#Write matches asynchronously, which may be faster for NFS.
param_asynchronous_write 0
#Limit the number of worker threads (experimental)
param_maximum_cpu_usage 0
#Flush stdout for each line of log when set to 1
param_stdout_line_flush 0
#When set to 1, the program will try to be deterministic by using 
#fixed seed for randomness, and disable threading in a few places
param_deterministic_behaviour 0

#Configuration version
param_config_version 52
```
