## Supplementary material for "Open source 3D phenotyping of chickpea plant architecture across plant development": Plantmodelbreakdown.R

```
library(Rvcg)
library(threed)
library(rgl)
library(dplyr)
library(tidyverse)
library(ggplot2)
library(png)
library(ggpubr)

# Return list of all .ply models in working directory, including in sub-folders
files<-list.files(recursive = T,pattern = ".ply")

# Process all .ply models to return breakdown .CSV file containing info on all triangles of each mesh and a visual summary of area by height
for (i in files){
Mesh<-vcgPlyRead(i,updateNormals = FALSE,clean = FALSE)
df<-as.data.frame(Mesh)

# Reorder data into new dataframe dfxyz
corner1<-subset.data.frame(df,vorder==1)
corner2<-subset.data.frame(df,vorder==2)
corner3<-subset.data.frame(df,vorder==3)

centres<-corner1[14:16]

corner1[7:19]<-NULL
corner1[2]<-NULL
corner2[7:19]<-NULL
corner2[2]<-NULL
corner3[7:19]<-NULL
corner3[2]<-NULL

corner1$x1<-corner1$x
corner1$y1<-corner1$y
corner1$z1<-corner1$z
corner1[3:5]<-NULL

corner2$x2<-corner2$x
corner2$y2<-corner2$y
corner2$z2<-corner2$z
corner2[1:5]<-NULL

corner3$x3<-corner3$x
corner3$y3<-corner3$y
corner3$z3<-corner3$z
corner3[1:5]<-NULL

dfxyz<-cbind.data.frame(corner1,corner2,corner3,centres)

#Calculate side lengths and areas of triangles
dfxyz$A<- sqrt((dfxyz$x1 - dfxyz$x2)^2 + (dfxyz$y1 - dfxyz$y2)^2 + (dfxyz$z1 - dfxyz$z2)^2)
dfxyz$B<- sqrt((dfxyz$x1 - dfxyz$x3)^2 + (dfxyz$y1 - dfxyz$y3)^2 + (dfxyz$z1 - dfxyz$z3)^2)
dfxyz$C<- sqrt((dfxyz$x2 - dfxyz$x3)^2 + (dfxyz$y2 - dfxyz$y3)^2 + (dfxyz$z2 - dfxyz$z3)^2)
dfxyz$S<- sqrt(((dfxyz$A + dfxyz$B + dfxyz$C)/2) * ((dfxyz$A + dfxyz$B + dfxyz$C)/2 -dfxyz$A) * ((dfxyz$A + dfxyz$B + dfxyz$C)/2 -dfxyz$B) *((dfxyz$A + dfxyz$B + dfxyz$C)/2 -dfxyz$C))
dfxyz$S<- dfxyz$S/2

# Height from base of plant
dfxyz$h<-dfxyz$fcz - min(dfxyz$fcz)
dfxyz<-dfxyz[order(dfxyz$h),]

# Normalised height from base of plant
dfxyz$hn<-dfxyz$h/max(dfxyz$h)

# Cumulative area from base of plant
dfxyz$Scum<-cumsum(dfxyz$S)

# Normalised cumulative area
dfxyz$Scumn<-dfxyz$Scum/max(dfxyz$Scum)

# Save CSV
name<-gsub("Meshed/","", i)
name<-gsub(".ply","", name)
write.csv(dfxyz,str_c(name,".csv"))

#Graphical summary
#Round to 1 mm bins
dfxyz$roundedheight<-round(dfxyz$h,digits = 1)
dfround<-dfxyz %>% distinct(roundedheight,.keep_all = TRUE)
dfround$Smmheight<-c(dfround$Scum[1],diff(dfround$Scum))

Areagraph <- ggplot(dfround,aes(x=roundedheight,y=Smmheight))+
  coord_flip()+
  # stat_smooth(se=FALSE,span=0.25,size=1,alpha=0.25)+
  geom_histogram(stat = "bin2d",bins = nrow(dfround))+
  labs(x=expression(paste("Height from base of plant /cm")),y=expression(paste("Area per 1mm height /cm"^2)))+
  labs(color="Genotype")+
  # scale_fill_brewer(palette = "Set1")+
  # scale_colour_brewer(palette = "Set1")+
  theme(plot.title = element_text(hjust = 0.5),
        axis.line = element_line(linetype = "blank"),
        panel.border = element_rect(colour = "black", fill=NA, size=1),
        panel.grid = element_blank(),
        panel.background = element_rect(fill = "gray90"),
        axis.ticks.y = element_blank(),
        axis.text.y=element_text(size=12),
        axis.text.x=element_text(size=12),
        axis.title=element_text(size=15),
        legend.key = element_rect(fill = "grey", color = NA),
        legend.text = element_text(size=12),
        legend.title = element_text(size=12,face = "bold"),
        legend.position = "none",
        legend.title.align = 0.5,
        strip.text.x = element_text(size = 14),
        text=element_text(family="Helvetica"))


plot3d(Mesh,xlab="",ylab="",zlab = "", axes=FALSE,main=NULL,sub=NULL,aspect = TRUE,lit=TRUE,alpha=0.5,col="#008000")
rgl.viewpoint( theta = 180, phi = 90, fov = 0, zoom = 0.65)
rgl.bringtotop()
par3d(windowRect = c(40, 40, 1080, 1080))
rgl.snapshot("image.png",fmt = "png")
image<-readPNG("image.png")
imagegg<- ggplot()+
  background_image(image)
imagegg

outputfig<-ggarrange(Areagraph, imagegg, ncol = 2, nrow = 1)
outputfig
ggsave(str_c(name,".pdf"), width = 12, height = 6)
rgl.clear()
}
```
