## Supplementary material for "Open source 3D phenotyping of chickpea plant architecture across plant development": VisualSFMbatch.bat

```
rem update filepath to folder containing VisualSFM exectutable
cd C:\Users\USERNAME\Documents\VisualSFM_windows_cuda_64bit

rem update filepaths to image sequence folders and output folders below, add more if needed, processing will proceed sequentially.
start /wait cmd /c VisualSFM sfm+pmvs C:\Users\USERNAME\Documents\3D_Scanner_Image_Folder\Plant1 C:\Users\USERNAME\Documents\3D_Scanner_Image_Folder\UncleanedPointClouds\Plant1.nvm
start /wait cmd /c VisualSFM sfm+pmvs C:\Users\USERNAME\Documents\3D_Scanner_Image_Folder\Plant2 C:\Users\USERNAME\Documents\3D_Scanner_Image_Folder\UncleanedPointClouds\Plant2.nvm
start /wait cmd /c VisualSFM sfm+pmvs C:\Users\USERNAME\Documents\3D_Scanner_Image_Folder\Plant3 C:\Users\USERNAME\Documents\3D_Scanner_Image_Folder\UncleanedPointClouds\Plant3.nvm
start /wait cmd /c VisualSFM sfm+pmvs C:\Users\USERNAME\Documents\3D_Scanner_Image_Folder\Plant4 C:\Users\USERNAME\Documents\3D_Scanner_Image_Folder\UncleanedPointClouds\Plant4.nvm
start /wait cmd /c VisualSFM sfm+pmvs C:\Users\USERNAME\Documents\3D_Scanner_Image_Folder\Plant5 C:\Users\USERNAME\Documents\3D_Scanner_Image_Folder\UncleanedPointClouds\Plant5.nvm
```
